# An *in utero* exposure to the synthetic estrogen diethylstilbestrol affects the fat pad composition in post-natal mammary glands

**DOI:** 10.1101/2025.09.11.675719

**Authors:** David Tovar-Parra, Alec McDermott, Jysiane Cardot, Melany Juarez, Fabien Joao, Rhizlane El Omri-Charai, Line Berthiaume, Bhawna Dhawan, Arash Aghigh, Yann Breton, François Légaré, Géraldine Delbès, Martin Pelletier, Étienne Audet-Walsh, Isabelle Plante

## Abstract

*In utero* exposure to the synthetic estrogen diethylstilbestrol (DES) has been linked to developmental abnormalities and elevated breast cancer risk in adulthood in human and rodent models. While the impact of DES on the mammary epithelium has been thoroughly investigated, its effect on the other cell types of the mammary gland remains understudied. Here, given that the mammary gland development is strongly associated with its microenvironment, we aimed to investigate how *in utero* DES exposure alters the mammary gland’s stromal and immune function across key developmental stages. To achieve this aim, timed-pregnant rats were gavaged daily with DES or vehicle from gestation days 16-21, and female offspring mammary glands were analyzed at pre-puberty (postnatal day 21 (PDN21)), puberty (PND46), and adulthood (PND90). We assessed morphological and extracellular matrix changes, performed transcriptomic cell-type enrichment analysis, measured cytokine expression, and quantified immune cell populations. DES-exposed mammary glands exhibited pronounced stromal remodeling, including increased collagen deposition and orientation by adulthood. Gene expression profiling indicated DES-induced stage-specific immune alterations: immune cell signatures were enriched at PND21 and PND90 but diminished at PND46. Correspondingly, DES increased macrophage populations at PND21 while reducing T-lymphocyte numbers at PND46 and PND90. DES exposure also dysregulated inflammatory cytokine/chemokine expression in adult glands, suggesting a persistent inflammatory environment. In conclusion, *in utero* exposure to an estrogenic compound can reprogram mammary development, inducing long-term changes in the extracellular matrix and immune landscape. These disruptions to stromal-immune homeostasis may impair normal mammary morphogenesis and increase susceptibility to breast pathologies later in life.

## Introduction

The development of the mammary gland is highly dynamic (Tovar-Parra et al. 2025), and is orchestrated by the regulation and interaction between epithelial cells, stromal components, and systemic hormonal signals (Biswas et al. 2022; Monkkonen et al. 2017). The transition between the pre-puberty, peri-puberty, and adulthood stages is characterized by processes such as ductal elongation, branching morphogenesis, adipocyte differentiation, extracellular matrix (ECM) remodeling, and immune cell infiltration (Andrechek et al. 2008; Need et al. 2014; Stewart et al. 2019). These morphological changes are regulated by hormones, principally by estradiol signaling through binding to its nuclear estrogen receptors (ERs) (Gruber et al. 2002; Li et al. 2010; Straub 2007). Among others, ERs trigger the transcription of mediators such as activator protein 1 (AP-1), nuclear factor kappa B (NF-*κ* B), signal transducer and activator of transcription 3 (STAT3), and stimulating protein 1 (SP1), which in turn regulate genes involved in epithelial and stromal cell proliferation, survival, migration, and metabolic regulation (Brisken and O’Malley 2010; Cheskis et al. 2007; Lacouture et al. 2023) Although the fetal development of the mammary gland is considered independent of hormones, however, ERα expression has been detected perinatally in humans (Keeling et al. 2000; Naccarato et al. 2000) and rodents (HoveyTrott and Vonderhaar 2002). In addition, it has been suggested that maternal hormones have a programming function in various tissues, enabling the actions of hormones later in life (Berenbaum and Beltz 2011; Wallen and Hassett 2009).

Our recent data indicate that the mammary gland development undergoes significant changes during the transition from pre-puberty to peri-puberty (Tovar-Parra et al. 2025). Over 1,500 genes exhibited substantial alterations, with pathways related to proliferation, lipid metabolism, and immune response being particularly impacted. Notably, there is a pronounced decrease in total fatty acid concentration during this transition, likely attributed to the downregulation of lipid metabolism genes such as fatty acid synthase (*Fasn)* (Tovar-Parra et al. 2025). Consistently, disruption of these physiological pathways by exposure to exogenous estrogens, estrogen-mimicking endocrine disruptors, or estrogen deficiency has been linked to long-term alterations in tissue composition, increased breast cancer risk, and impaired lactational capacity (Brisken and Ataca 2015; Makris 2011; Mallepell et al. 2006).

Diethylstilbestrol (DES) is a xenoestrogen compound with high ERα affinity (Al Jishi and Sergi 2017; Fenton 2006; Soto and Sonnenschein 2015). Pregnant women were prescribed DES to prevent complications of pregnancy (Al Jishi and Sergi 2017; Ervin Adam et al. 1976). However, its use during pregnancy has been associated with increased risk of breast cancer for both the mothers and their daughters, as well as with different pathologies as clear cell adenocarcinoma, reproductive tract abnormalities, and fertility problems (Hilakivi-Clarke 2014; Singh et al. 2015). Mechanistically, DES has been linked with modulations of DNA methylation, gene expression, and histone modifications (Bredfeldt et al. 2010). In rats, neonatal exposure (at birth) to a single subcutaneous dose of 1 µg of DES resulted in an increased number of terminal end buds (TEBs) and highly proliferative structures associated with mammary ductal elongation, at postnatal day (PND) 50 compared to controls (Ninomiya et al. 2007). Furthermore, a similar exposure resulted in increased TEBs at PND35, as well as increased mRNA expression of β-casein and whey acidic protein, genes related to differentiation of terminal duct lobular units (TDLUs) and milk production after pregnancy (Umekita et al. 2011). Although previous studies have documented DES-exposure changes in ductal morphology and hormone receptor expression (Hovey et al. 2005), the impact on the mammary stroma, especially on adipocyte, lipid metabolism, immune cell populations, cytokine and chemokine profiles and ECM organization remains poorly understood.

In this study, we hypothesized that *in utero* exposure to DES during late gestation induces stage-dependent reprogramming in the cellular and molecular mechanisms of mammary gland development, manifesting as changes in epithelial architecture, stromal adipogenesis, lipidomic distribution, immune cell composition and signaling, and ECM structure across developmental stages. Furthermore, the effects on the stroma should be more critical when rats are exposed from late gestation, a window of exposure that corresponds to the period when the fat pad is thought to begin its differentiation. To test this hypothesis, we employed an integrative, multi-omics approach combining transcriptomic, lipidomic, and immune proteomic analyses.

## Materials and methods

### Animal treatment

The protocol was approved by the Institutional Animal Care and Use Committee at the INRS (# 2202-02), and the animals were treated following the guidelines outlined by the Canadian Council on Animal Care. Female virgin Sprague-Dawley rats were exposed daily to DES (1 and 10 µg/kg of body weight (BW)/day) or corn oil containing vehicle (4%DMSO, vehicle control) via gavage from GD16 until GD21. The stock solutions were prepared using DES (Sigma-Aldrich D4628, purity ≥98%) at concentrations of 5 and 50 µg/mL dissolved in 100% dimethyl sulfoxide (DMSO) (Sigma-Aldrich, Cat. D8418, purity ≥99%). Working solutions corresponding to final concentrations of 1 and 10 µg/mL were prepared by 5-fold dilution of the stock solution in corn oil (vehicle control) for oral gavage. For the sample collection, rats were euthanized by CO_2_ inhalation followed by cervical dislocation at postnatal day 21 (PND21), corresponding to pre-puberty, PND46 (peri-puberty), and PND90 (adulthood). For each group, 5-7 animals were euthanized, and four pairs of mammary glands (two thoracic, one abdominal, one inguinal pair) were taken.

### Whole mount analysis

For the epithelial tissue analysis, the right inguinal mammary gland for each animal was dissected and processed according to previously described protocols (Crobeddu et al. 2022; Gouesse et al. 2019; McDermott et al. 2025; Tovar-Parra et al. 2025). Briefly, mammary glands were spread on slides and fixed overnight at room temperature (RT) in Carnoy’s solution, composed of a 6:3:1 mixture of 100% ethanol, chloroform, and glacial acetic acid. Samples then underwent rehydration through a serial ethanol gradient from 70% to 0% and were stained overnight at RT in a carmine alum solution (2% carmine with 5% potassium aluminum sulfate in water), followed by dehydration in successive ethanol baths, clearing in xylene, and final mounting in Permount (Fisher Chemical, Ontario, Canada, no. SP15-500).

Digital images of the prepared slides were captured using a Canon PowerShot G9x camera on a transilluminator (Henning Graphics LR299343) with a measurement scale to ensure dimensional accuracy. For each group (N = 5–7 animals per group), the epithelial area was quantified from the whole-mount images by calibrating the scale and measuring the area in cm² using the skeletonize tool in ImageJ software (https://imagej.net/Fiji/Downloads). Branching complexity was assessed by selecting three to five representative sections from each skeletonized image and computing the number of intersections per cm² using the Neuroanatomy Sholl analysis plugin (StankoEasterling and Fenton 2015; Stanko and Fenton 2017).

### Histology

The thoracic mammary glands were dissected and immediately embedded in OCT Cryomatrix (Fisher Scientific, Cat. 23-730-571) on dry ice before storage at –80°C. The frozen tissues were sectioned into 5 µm slices using a cryostat, fixed in Bouin’s solution (Sigma-Aldrich, Cat. HT10132) overnight at RT and stained following Masson’s Trichrome protocol. Briefly, sections were sequentially immersed in Weigert’s iron hematoxylin (10 min), Biebrich scarlet-acid fuchsin (15 min), phosphomolybdic-phosphotungstic acid (20 min), and aniline blue (5 min) with water washes after each step and then treated with 1% acetic acid (5 min). Subsequent dehydration in 70, 95, and 100% ethanol (5 min each) and clearing in xylene (5 min) preceded final mounting with coverslips and Permount (Fisher Chemical, Ontario, Canada, No. SP15-500) (Plante Stewart and Laird 2011). Confocal images were acquired using a Nikon A1R+ microscope equipped with a digital camera using NIS-Element software.

ImageJ was used to analyze the images. Intact adipocyte areas were determined with the drawing tool, and overall adipose tissue area and cell counts were quantified using the Adiposoft plugin (Galarraga et al. 2012). Additionally, mammary gland component analysis was performed on these images by quantifying the proportions of epithelial tissue (red), collagen (blue), and adipose tissue (white) through the segmentation of unique pixel characteristics based on hue, saturation, and intensity using NIS-Element Nikon’s analysis software. This method generated distinct layers via threshold selection and object counting, thereby allowing precise determination of the relative components within each image (Crobeddu et al. 2022).

### Lipidomic analysis

The lipid profile was generated from snap-frozen inguinal mammary glands. Tissues were first pulverized in liquid nitrogen on dry ice and weighed before homogenization in 0.9% NaCl. The internal standard phosphatidylcholine C21:0 (Sigma-Aldrich) was added to each homogenate, and lipids were then extracted using a chloroform:methanol (2:1, v/v) solution according to a modified Folch method (Shaikh and Downar 1981). Following centrifugation, the lipid phase was collected and dried under a nitrogen stream. The dried extracts were subsequently resuspended in 2 mL of methanol/benzene (4:1, v/v) and methylated with acetyl chloride, following our standard protocol (Tovar-Parra et al. 2025). The resulting fatty acid profiles were analyzed using capillary gas chromatography with a temperature gradient on an HP5890 gas chromatograph (Hewlett Packard, Toronto, Canada), equipped with an HP-88 capillary column (100 m × 0.25 mm i.d. × 0.20 µm film thickness; Agilent Technologies) and a flame ionization detector, with helium as the carrier gas at a 50:1 split ratio. Fatty acids were identified based on their retention times by comparison with standard mixtures (Tovar-Parra et al. 2025). Data was expressed as a percentage of total fatty acids and as mg of FA/g of mammary gland tissue.

### Transcriptomic analysis

RNA was extracted from 100 mg of inguinal mammary gland tissue per animal from four randomly chosen samples per group using the Aurum Total RNA Fatty and Fibrous Tissue Kit (Bio-Rad, Ontario, Canada, Cat. 7326830) following the manufacturer’s instructions. Purity was determined by Nanodrop (Thermo Scientific), and RNA integrity and concentration were confirmed using the Agilent 2100 Bioanalyzer with the RNA 6000 Nano Kit, ensuring that all samples exhibited a RIN > 8. The extracted RNA samples were then sent to the Genomic Centre of the Centre de recherche du CHU de Québec – Université Laval for RNA sequencing on a HiSeq 2500 platform (125 bp paired-end sequencing). For sequencing data analyses, raw sequencing reads were initially quality-checked using FastQC v0.12.0, then trimmed with TrimGalore v0.6.10, and reassessed with FastQC before being aggregated with MultiQC v1.2. High-quality trimmed reads were pseudo-aligned to the Rattus norvegicus Rnor_6.0 reference transcriptome (https://may2021.archive.ensembl.org/Rattus_norvegicus) using Kallisto v0.46.1, and the resulting data were deposited in NCBI under accession number PRJNA1250603.

Differential gene expression analysis was conducted using the DESeq2 v1.42.1 package in R, with significant genes defined by a Benjamini-Hochberg adjusted p-value <0.05 and an absolute log2 fold change ≥1.5. K-means clustering was applied to determine molecular signatures and average expression levels across groups, while Gene Set Enrichment Analysis (GSEA) was used to explore regulated pathways and cell type signatures (Kanehisa et al. 2025). Further biological insights were obtained using Gene Ontology (GO) and KEGG enrichment tools via the SRplot package (http://www.bioinformatics.com.cn) and the R software environment, and enriched pathways and term similarities were visualized with Metascape (https://metascape.org/) and Enrichr (https://maayanlab.cloud/Enrichr/).

### Second harmonic generation microscopy

A custom-built second harmonic generation (SHG) microscope was employed to capture signals from the 5 µm tissue cryosections. The light source was a Ytterbium fiber laser (MPB Communications Inc., Montréal, CA) featuring a 125-fs pulse duration, a wavelength of 1030 nm, and a 25 MHz repetition rate, delivering up to 3 W of average power. To avoid thermal damage, a power control unit was used, comprising a half-wave plate and a Glan-Thompson polarizer, which regulated the output power, was adjusted between 30 mW and 50 mW (corresponding to pulse energies of 1.2 to 2 nJ at the objective focus), depending on the sample. Illumination was achieved using an air objective (UplanSApo, Olympus, Japan) with 10× magnification and a numerical aperture of 0.3, while the sample was scanned over an average area of 6000×6000 µm² using a high-speed motorized XY stage (MLS203; Newton, NJ, USA).

The distance between the sample and the objective was fine-tuned along the z-axis through a combination of mechanical and piezoelectric motors (PI Nano-Z, USA). A condenser collected both the fundamental and SHG signals, and a multi-stage filtration process, incorporating two short-pass filters (Semrock, 720 nm blocking edge) and a 515/30 nm band-pass filter (Semrock), was used to isolate the SHG signal from the fundamental laser light. Signal detection was performed by a photomultiplier tube (R6357, Hamamatsu Photonics) set to 800 V, with synchronization managed via a multichannel I/O board (National Instruments) and a custom Python program that allowed real-time adjustment of the imaging area and resolution (Pinsard 2020). Raw images were subsequently visualized using Fiji-ImageJ software.

In addition, a polarization SHG (P-SHG) study was conducted on the mammary gland samples to assess collagen fiber alignment. Each sample was imaged using linearly polarized light at varying azimuthal angles (Ω) adjusted by a motorized half-wave plate inserted in the laser path (Ducourthial et al. 2019; Golaraei et al. 2020). A series of 18 images, acquired in 20° increments from 0° to 340°, provided the data for analysis. Collagen fiber alignments (θ) were computed from these intensity images as a function of the incident light angle using the relationship:

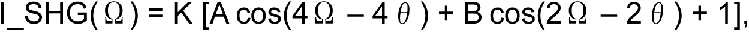

where K represents the mean photon count per image, and A and B are susceptibility constants. Further details regarding the algorithm and procedural specifics are provided in the referenced literature (Aghigh et al. 2024; Teulon et al. 2015).

### Immunofluorescence

Thoracic mammary gland slides were analyzed using immunofluorescence staining, following methods previously described (Gouesse et al. 2019). Briefly, 5 µm tissue cryosections were fixed in 4% formaldehyde and subsequently blocked in 3% BSA diluted in PBS. The sections were then incubated with a primary antibody (Supplementary Table 1). Primary antibodies were diluted in 3% BSA-PBS for 60 minutes at RT. After three 5-minute washes, cryosetions were incubated with the corresponding secondary antibody for 60 minutes at RT (Supplementary Table 1). After three 5-minute washes, the sections were stained with 4′, 6-diamidino-2-phenylindole (DAPI, 62248 – Thermo Scientific) and mounted using Fluoromount-G (Cedarlane, Burlington, Ontario, Canada). Immunofluorescence images were acquired with a Nikon A1R+ confocal microscope equipped with a spectral detector, and all subsequent analyses were performed using Fiji-ImageJ software.

### Cytokine and chemokine quantification

Snap-frozen inguinal mammary glands were pulverized in liquid nitrogen on dry ice and weighed prior to protein extraction using the ProcartaPlex™ Cell Lysis Buffer Kit (Cat. N° EPX-99999-000, Thermo Fisher Scientific), following the manufacturer’s instructions. Protein concentration was determined using the Pierce BCA protein assay kit (Thermo Scientific, Rockford, Illinois, USA), and all samples were normalized to 10 mg/ml. Subsequently, cytokines and chemokines were quantified using the Cytokine & Chemokine 22-Plex Rat ProcartaPlex™ Panel (Cat. N° EPX220-30122-901, Thermo Fisher Scientific) on a Luminex platform, according to the manufacturer’s protocols.

In brief, the assay plate was first seeded with capture beads that were subsequently washed using a hand-held magnetic plate washer. A 25 µL buffer was added to each well, followed by the addition of 25 µL of sample per well and the standard curve. After designated incubation (90 min) and wash periods (3 times), a 25 µL biotinylated detection antibody mixture was added to the plate. Following further incubation (30 min) and washing, 50 µL of Streptavidin-PE was introduced, following incubation (30 min), and the plate was then analyzed on a Bio-Plex™ 200 system (Bio-Rad).

### Statistical analysis

All statistical analyses were performed using GraphPad Prism 9 and the RStudio environment. Initially, descriptive statistics, including mean, median, and standard deviation, were calculated to summarize the data. To assess the distribution of the data, we utilized both the Shapiro-Wilk and Kolmogorov-Smirnov normality tests. For datasets meeting the assumptions of normality, parametric tests were applied, including one-way ANOVA for multiple group comparisons and Student’s t-test for pairwise comparisons; for datasets that did not meet these assumptions, non-parametric alternatives such as the Kruskal-Wallis test and the Mann-Whitney U-test were employed. Pearson’s correlation coefficient was computed to evaluate the strength and direction of relationships between variables. For all statistical analyses, a p-value < 0.05 was considered indicative of a statistically significant difference.

## Results

### Impact of in utero DES exposure on epithelial mammary gland morphology

To investigate the morphological consequences of *in utero* exposure to DES, pregnant rats were treated with 1 or 10 µg/kg/day from GD16 until GD21 (Fig. S1). Mammary glands from rat offspring were evaluated at key developmental stages: pre-puberty (post-natal day 21 (PND21)), peri-puberty (PND46), and adulthood (PND90). No significant differences were found in rats’ weight treated with DES at 1 or 10 µg/kg/day compared with controls at any time point (Fig. 1A). In contrast, normalized mammary gland weight was significantly higher in the 10 µg/kg/day group compared with the control at PND46 (Fig. 1A). Whole-mount analysis of the right inguinal gland was done to quantify the epithelial development (Fig. 1B). No significant differences were detected in total epithelial area across any age or treatment group (Fig. 1C). However, exposure to 10 µg/kg/day of DES resulted in a significant reduction in epithelial branching complexity at PND90 compared to controls (Fig. 1C). All together suggest that exposure to the highest dose of DES at the end of gestation significantly increases mammary gland mass at peri-puberty and impairs epithelial branching complexity at adulthood, suggesting long-term effects of the treatment.

**Figure 1.**
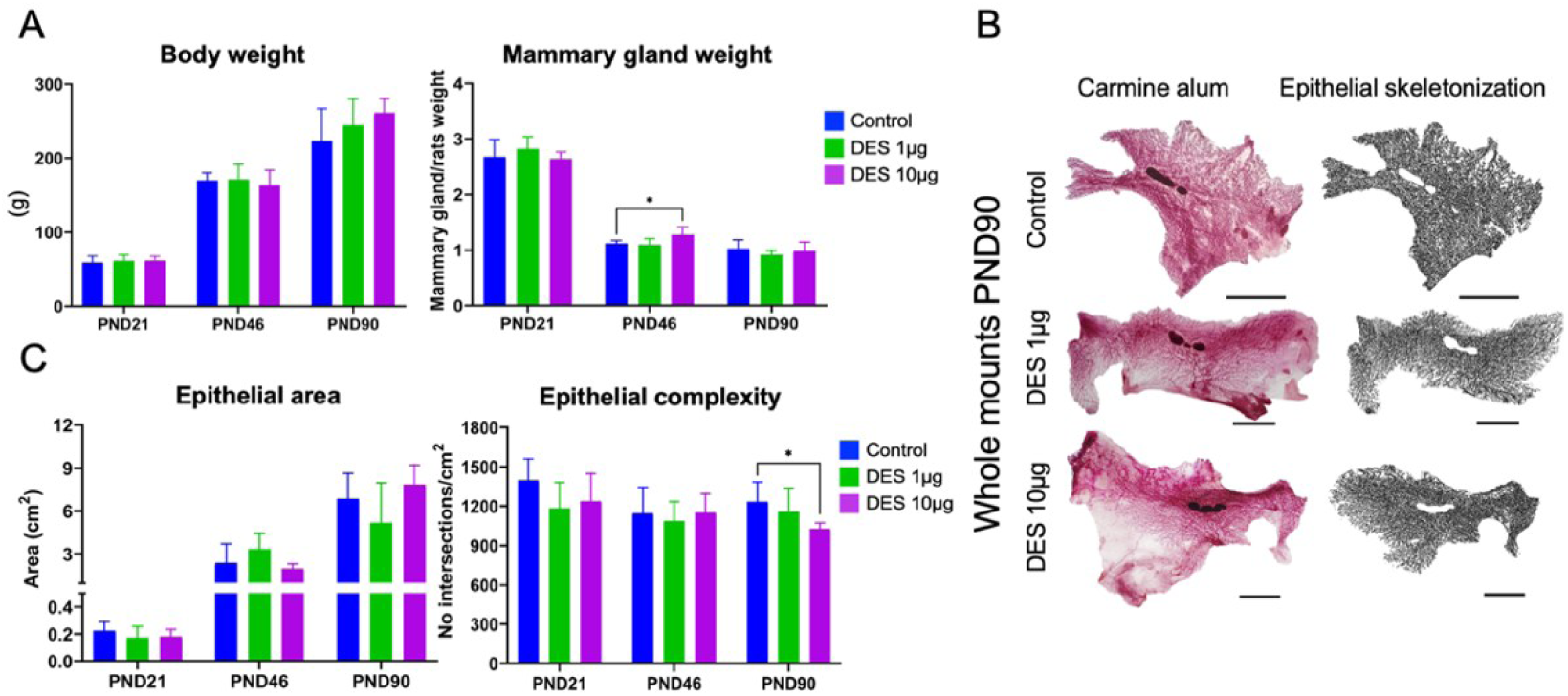
Modulatory effect of *in utero* exposure to DES (1 or 10µg/kg/day) on postnatal ductal development in rats. **A.** Average body and mammary gland weight normalized to total rat weight per group. **B.** Representative images of whole mount staining and skeletonized mammary gland epithelium obtained using ImageJ for each group at PND90 (scale bar 1 cm). **C.** Quantitative epithelial area and complexity for each condition on each stage obtained using whole-mount images. Graphs represent the average of the group (N ≥ 5) with SD (*p<0.05).

### Impact of in utero DES exposure on stromal mammary gland morphology

We then evaluated the effects of DES on the stroma, as the exposure period utilized for this study was chosen to specifically affect this compartment. We first determined the number and size of adipocytes using Masson’s Trichrome stain (Fig. 2A-C). A significant increased number of adipocytes was measured at the 1 µg/kg/day dose at PND46 and PND90 (Fig. 2D). The number of adipocytes was also significantly increased for the 10 µg/kg/day dose at both PND21 and PND90 (Fig. 2D). At PND90, the mean size of adipocyte was significantly larger in the 10 µg/kg/day group compared with controls (Fig. 2E). Finaly, we used Nikon’s segmentation software to quantify the proportion of the epithelial (red), collagen (blue), and adipose (white) compartments in each mammary gland section (Fig. S2A-D). No changes were observed in collagen and epithelial content across all groups (Fig. 2F-G). However, exposure to 1 µg/kg/day DES led to a significant decrease in adipocyte proportion at PND21 and PND46 (Fig. 2H). These results suggest that exposure to both doses of DES at the end of gestation, significantly affects the number and proportion of adipocytes but has little impact on their size in pre-puberty, peri-puberty and adulthood.

**Figure 2.**
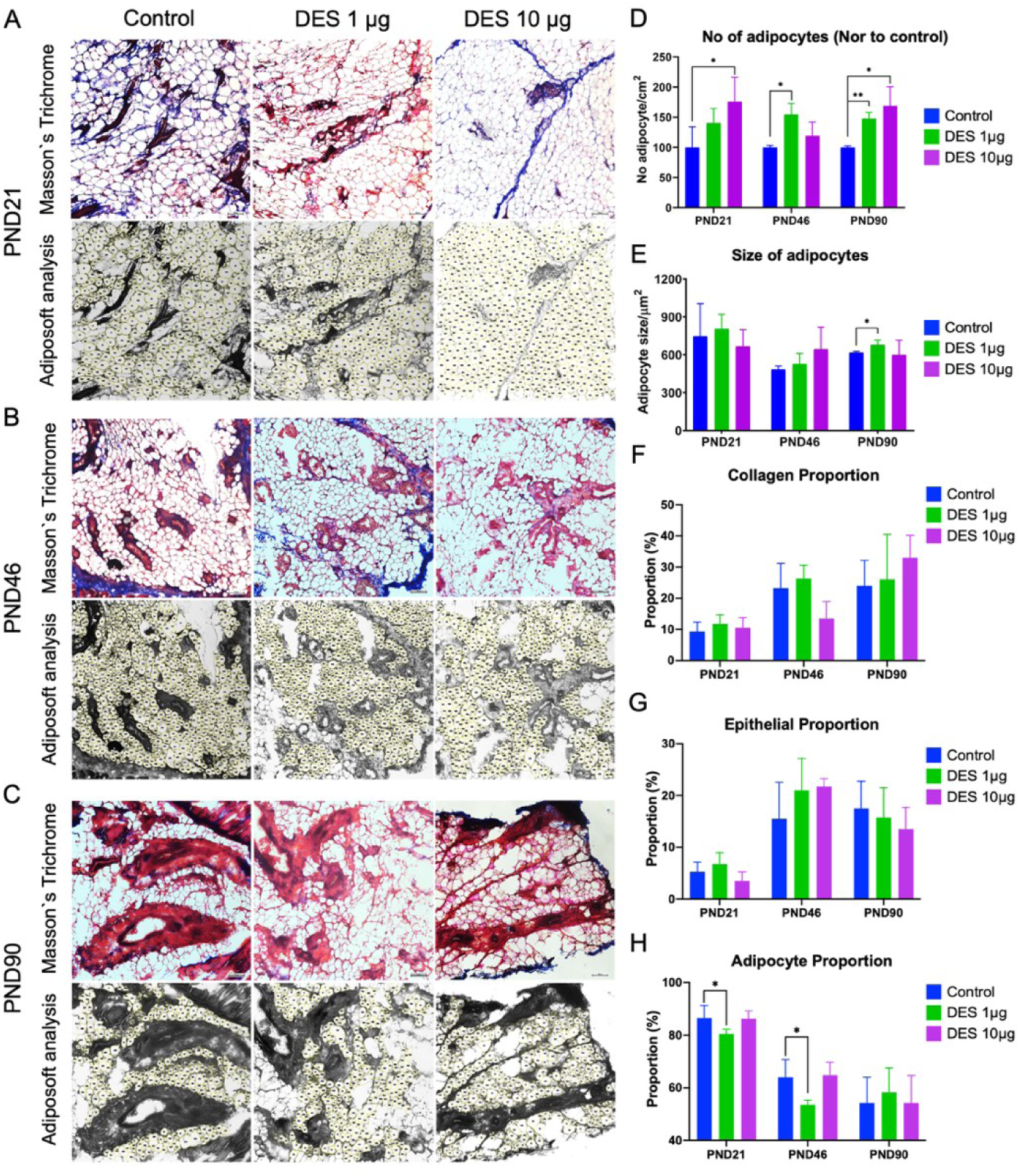
Morphological changes in adipocytes and proportion of mammary gland compartments across developmental stages after DES exposure (1 or 10 μg/kg/day). Representative Masson’s Trichrome-stained images of mammary gland sections from PND21 (**A**), PND46 (**B**), and PND90 (**C**) control rats and rats exposed to DES at 1-10 μg/kg/day. Scale bar 100 μm. (D-E) Histograms representing the mean number and size of adipocytes per cm² and μm², respectively. (F-H) Tissue proportions include collagen, epithelium, and adipocytes. For all graphs, error bars represent SD (*p < 0.05, **p < 0.001).

### Alteration in the mammary gland fatty acid profiles after in utero exposure to DES

To investigate whether changes in adipocyte size and number correlate with shifts in lipid metabolism, we performed targeted lipidomic profiling of the mammary glands. We quantified a total of 43 fatty acids that were further classified into 15 categories according to fatty acid characteristics (Table 1). Our analyses showed that 10 out of 43 detectable fatty acids were deregulated across developmental stages in the 1 µg/kg/day group, whereas only 3 fatty acids changed significantly in the 10 µg/kg/day group (Table 1).

**Table 1.**
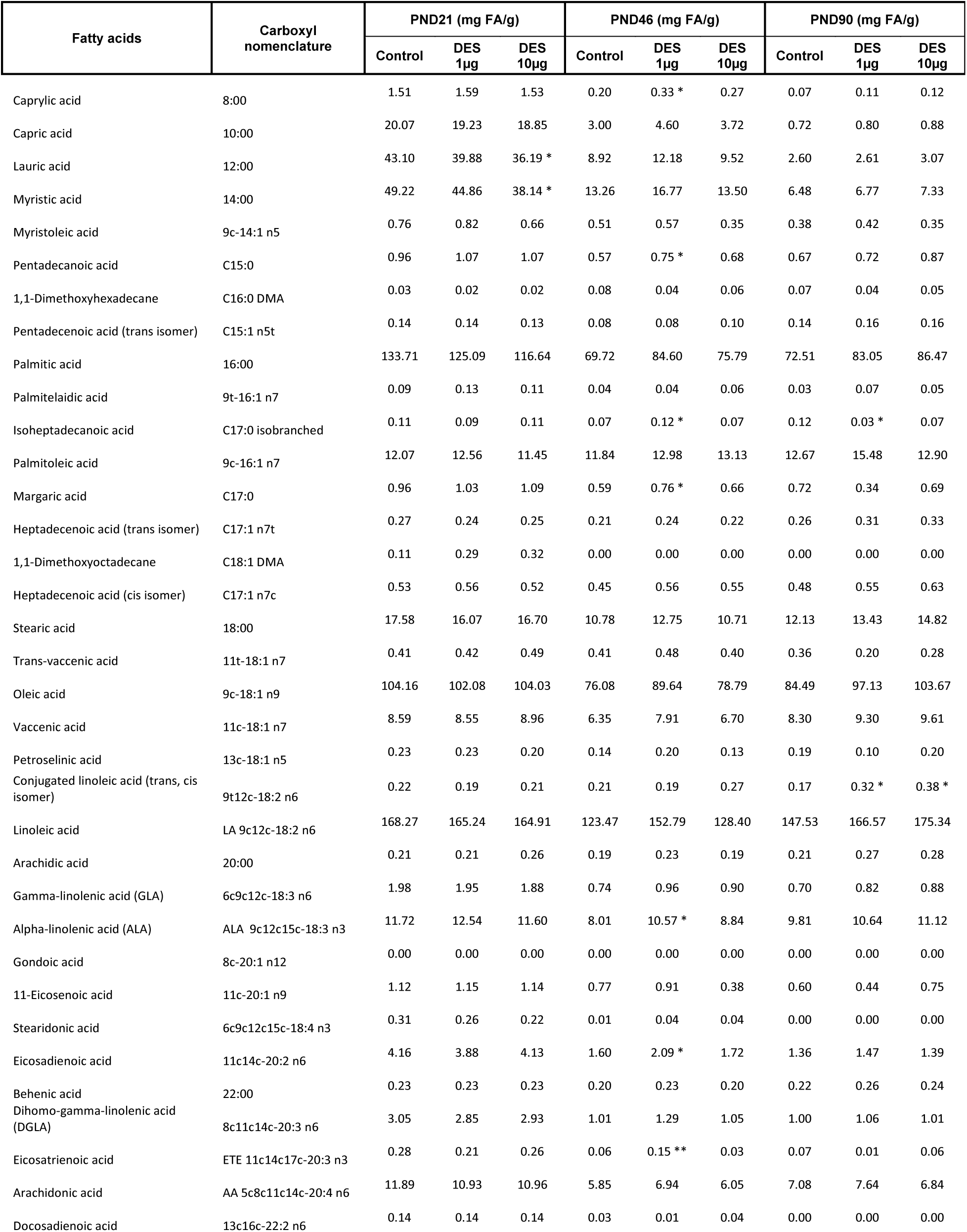
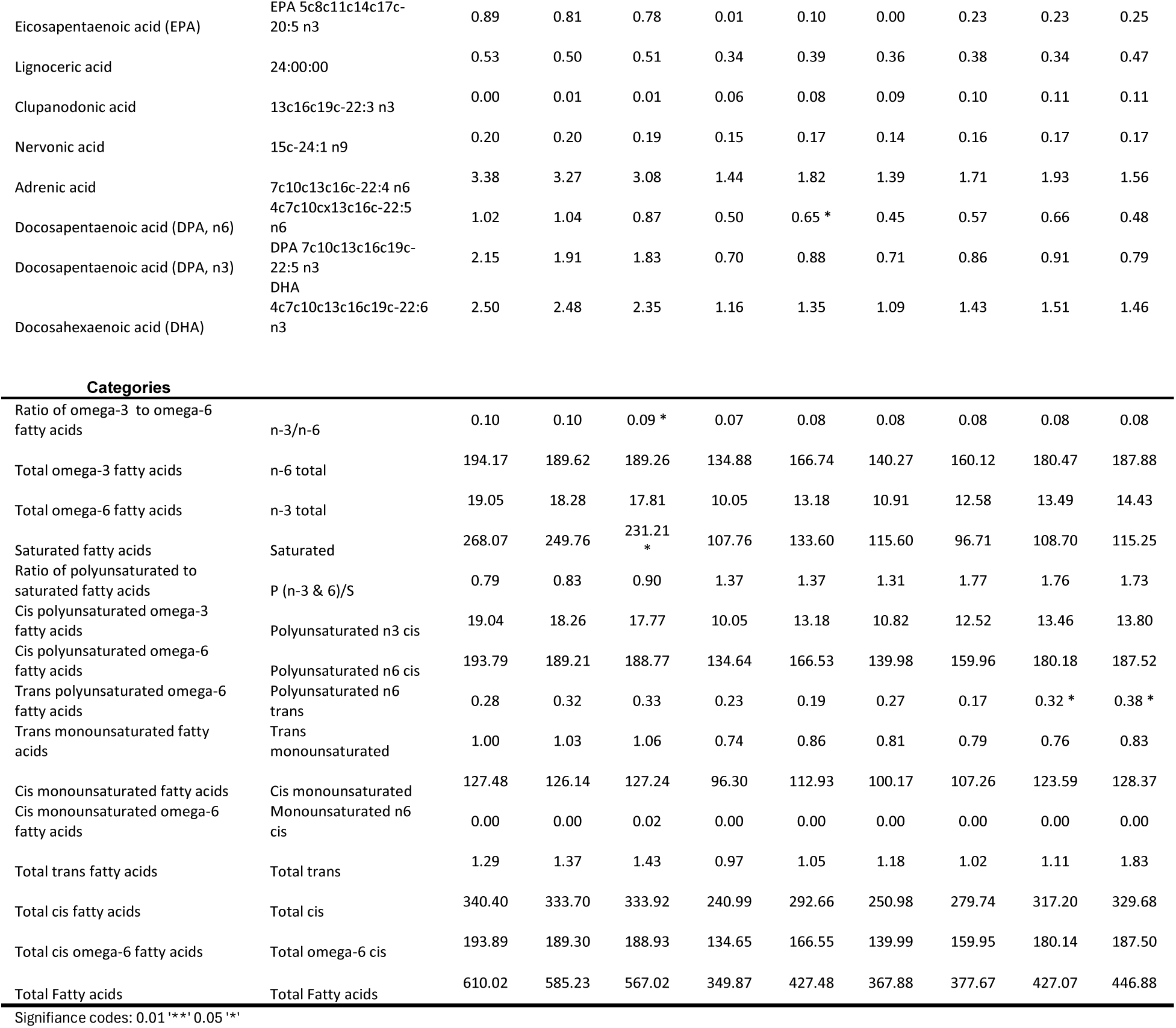
Fatty acid profiles in mammary glands across developmental stages and DES exposure conditions.

In the Pearson correlation analysis, we observed more important global distribution changes at 1 µg/kg/day of DES at PND46 and PND90 compared to the control, then at PND21 and the 10 µg/kg/day dose of DES (Fig. 3A, and Fig. S3A-B). Interestingly, at PND21, the animals treated with the highest dose of DES exhibited a reduction in saturated fatty acids such as lauric (c12:0) and myristic (c14:0) acids compared to controls (Fig. 3B). In contrast, at PND46 and PND90, the 1 µg/kg/day group showed marked elevation in medium-chain (caprylic c8:0, pentadecanoic c15:0) and long-chain saturated species (margaric c17:0, isoheptadecanoic iso-c17:0) (Fig. 3B), as well as various polyunsaturated fatty acids (PUFAs) including *α* -linolenic (c18:3 n-3), eicosatrienoic (c20:3), docosapentaenoic (c22:5 n-3), linoleic (c18:2), and n-6 trans-PUFAs, which are implicated in membrane architecture, signaling, proliferation, and immune regulation (Fig. S3C) (Hadley et al. 2017; Ip et al. 1994; Reiss et al. 2011; Vara-Messler et al. 2017). In summary, these findings suggest that while the higher dose mainly decreases specific saturated fatty acids at the pre-pubertal stages, *in utero* exposure to DES at 1 µg/kg/day appears to permanently reconfigure mammary gland fatty acid metabolism, as evidenced by an increase in both saturated and unsaturated fatty acids during peri-puberty and adulthood.

**Figure 3.**
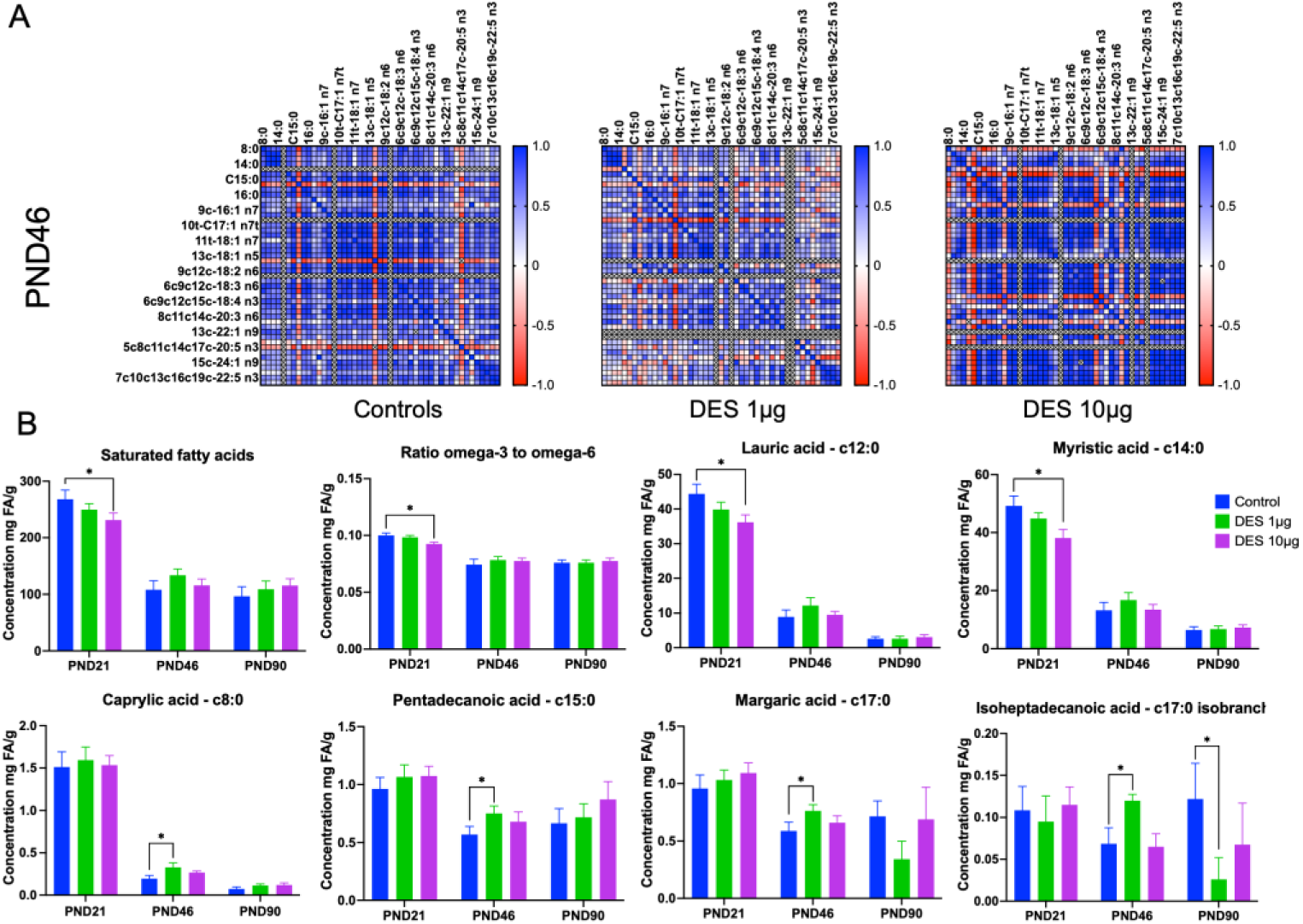
An *in utero* exposure to DES affects the lipid distribution in the mammary gland across developmental stages. **A.** Correlation of fatty acids showing the different lipid profiles at PND46; red indicates a negative correlation in the interaction between lipids, and blue indicates a positive correlation. **B.** Histogram representation of mean concentrations of saturated fatty acids, ratio of omega-3 to omega-6 fatty acids, and lauric, myristic, caprylic, pentadecanoic, margaric, and isoheptadecanoid fatty acids. Graphs represent the mean of the group with SD (*p < 0.05).

### Developmental stage-dependent remodeling of mammary extracellular matrix protein by in utero exposure to DES

Given the observed alteration in stromal components induced by *in utero* DES exposure, such as in adipocyte metrics and fatty acid profile, we analyzed other components of the stroma as the ECM. Immunofluorescence staining for collagen I, III, V, and laminin was thus performed on mammary gland sections collected at PND21, PND46, and PND90. At PND21, exposure to DES led to a reduction in collagens, which was significant for collagen I and V for the 1 μg/kg/day and both doses of DES, respectively (Fig. 4A, D and Fig. S4). In contrast, at PND46, the 10 µg/kg/day group exhibited a marked increase in both collagen I and laminin signal, while collagen III and collagen V levels remained unchanged (Fig. 4B, D and Fig. S5). By PND90, DES exposure had no significant effects on collagen I, III, V, or laminin levels (Fig. 4C, D and Fig. S6). These findings demonstrate that *in utero* DES exposure produces a dynamic, dose- and developmental stage-specific reorganization of the mammary ECM characterized by early depletion of fibrillar collagen components, a transient peri-pubertal surge in both fibrillar and basement-membrane proteins, and eventual normalization of matrix composition by adulthood.

**Figure 4.**
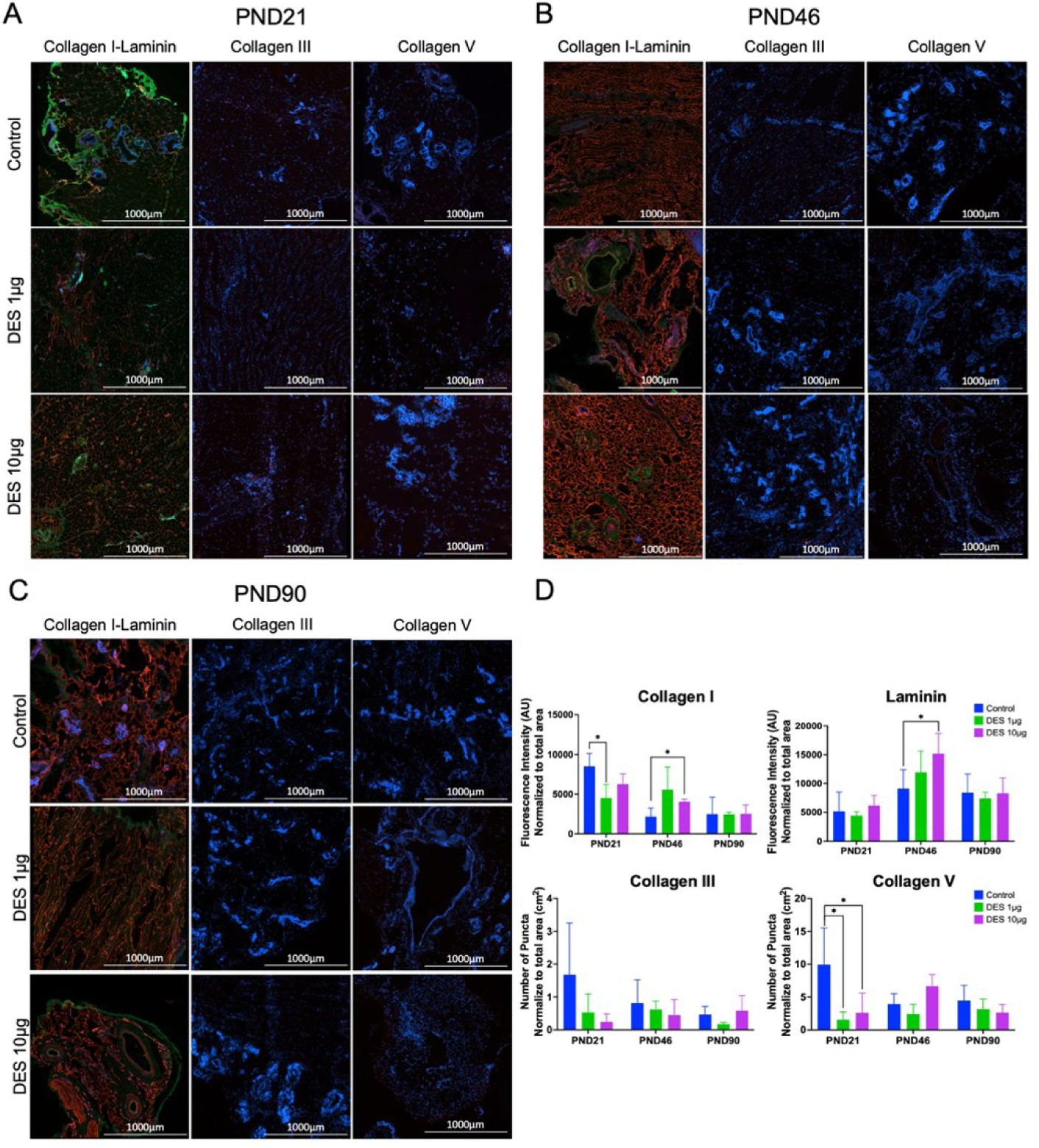
*In utero* DES exposure alters the collagen and extracellular matrix components in the stromal tissue of the mammary gland. Representative immunofluorescence images showing Collagen I (red) and laminin (green), Collagen III (red), and Collagen V (red) at PND21 (**A**), PND46 (**B**), and PND90 (**C**) in control and DES-treated groups (scale bar = 1000 μm). **D.** Histogram of fluorescence intensity quantifying collagen I, laminin, and normalized points per cm² for Collagen III and V in DES 1 and 10 μg/kg/day treatment groups. Graphs represent the mean of the group with SD (*p < 0.05).

### Collagen fiber remodeling in DES-exposed mammary glands

Collagen fibers are thought to guide epithelial ductal elongation during puberty, which aligns with pre-existing collagen fibers (Nerger et al. 2021). Building on observed alterations in collagens at PND21 and PND46 (Fig. 4), we further investigated the effects of DES on collagen fibers using conventional SHG and polarization-resolved SHG (p-SHG) with CurveAlign analysis on sections of mammary glands to comprehensively assess fibrillar architecture from pre-puberty through adulthood (Aghigh et al. 2024; Tilbury et al. 2014). P-SHG images were captured at 18 polarization states from 0 to 340 degrees in 20-degree increments, and fiber orientations were quantified in angle bins from −90 to 90 degrees. This binning is an equivalent representation of axial orientations modulo 180 degrees, so bins below 0° correspond to their 180-degree complements, in a 0-to-180-degree view, for example −30° equals 150° (Fig. 5A). At PND21, PND46, and PND90, no significant differences were observed. At PND21, a high inter-sample variability in the levels of fibers can be observed for both DES-treated group (Fig. 5B), with more fibers in negative angles, (correspond to their complements in the 0° to 180° representation) (Fig. 5C).

**Figure 5.**
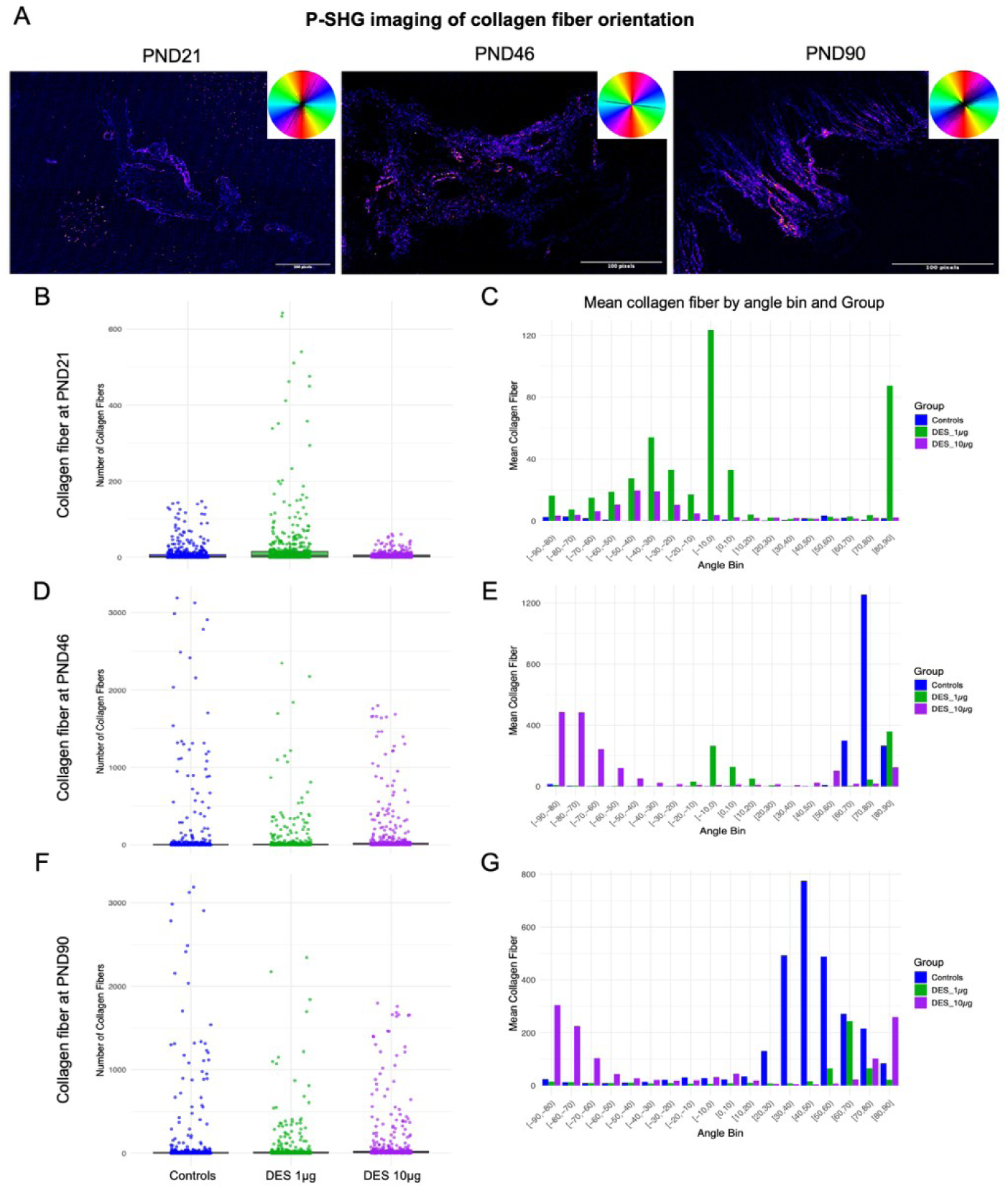
Effect on collagen content and fibers orientation after DES exposure in mammary glands assessed by second harmonic generation imaging. **A.** SHG image of collagen distribution with polar orientation plots at PND21, PND46, and PND90. Collagen fibers quantified at PND21 (**B**), PND46 (**D**), and PND90 (**F**) in the mammary gland of control rats and rats exposed to DES. **C.** Boxplot showing collagen concentration across orientation bins from −90 to 90 degrees (for equivalent range of 0 to 180 degrees) in 10 degree intervals for control, DES 1 μg per kg per day, and DES 10 μg per kg per day groups at PND21 (**C**), PND46 (**E**), and PND90 (**G**). All fiber orientations are axial and reported modulo 180 degrees. For display, bins are labeled from −90 to 90 degrees, where negative values are simply 180 degree complements of positive values, for example, −30 equals 150.

At PND46, variability in fiber counts was similar between groups (Fig. 5D), but orientations differed: controls were enriched between 60 and 90 degrees, the lower DES dose spanned 0 to 90 degrees with an additional mode near −10 degrees (170°), and the higher dose shifted toward −90 to −30 degrees (90 to 150 degrees) (Fig. 5E). At PND90, both DES groups tended to show fewer fibers overall (Fig. 5F), with the higher dose enriched in bins below 0 degrees (from 90 to 180 degrees) and controls and the lower dose enriched above 0 degrees (0 to 90 degrees) (Fig. 5G). While not statistically significant, these results suggest that collagen fibers are changing in orientation, which could potentially be associated with the lower complexity of the epithelium at PND90. Together, these data demonstrate that *in utero* exposure to DES induces subtle, yet persistent remodeling of collagen architecture marked by developmental increases in fiber density and developmental stage angular realignment of fibrils, thus contributing to the altered stromal microenvironment in DES-exposed mammary glands.

### Developmental stage-specific transcriptomic remodeling of the mammary gland following in utero DES exposure

To further understand mechanisms involved in structural and lipidomic alterations induced by *in utero* exposure to DES (1 µg/kg/day), we performed RNA-seq on the mammary gland at PND21, PND46, and PND90. The lowest dose of DES was chosen for those analysis as it resulted in greater lipidomic alterations at PND46 and PND90 (Fig. 3). Consistently, following pseudo/alignment (Kallisto) and rigorous quality controls (FastQC, MultiQC), to explore these transcriptional differences, we performed unsupervised k-means clustering (k=10) of normalized gene counts and identified clusters with distinct expression patterns (Fig. 6A). We found that in cluster 1, the relative gene expression was higher in the DES-exposed group across all stages (Fig. 6A). A pathway enrichment analysis of cluster 1 revealed that these genes were mainly associated with cell morphogenesis, cell growth and proliferation, and columnar/cuboidal differentiation (Fig. 6B). These gene signatures are consistent with changes in gland architecture, such as epithelial cell growth and differentiation, ductal elongation, branching formation, and TEB formation (Biswas et al. 2022). Genes from cluster 5 showed a low relative expression in the DES group compared to the control, specifically at PND46 (Fig. 6A). This cluster was enriched for lymphocyte activation, positive regulation of immune response, cytokine production, and innate immune response, among others (Fig. 6B).

**Figure 6.**
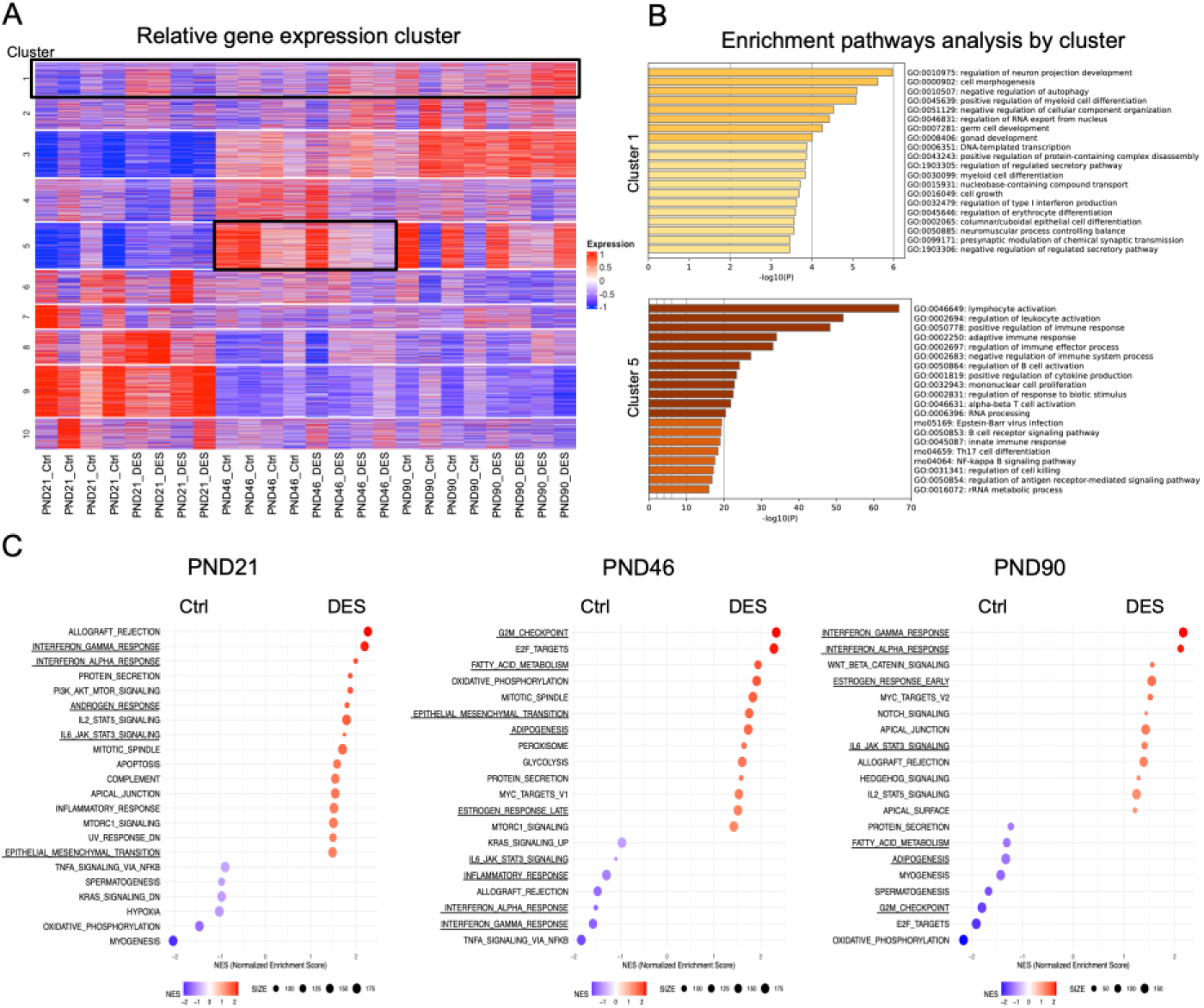
*In utero* exposure to 1 ug/kg/day of DES affects the transcriptional profiling and gene set enrichment analysis in mammary gland development. **A.** Heatmap represents gene expression clusters across all ages and treatment groups, with clustering performed using k-means (10 clusters identified). Clusters 1 and 5 are highlighted by black frames. Metascape enrichment network visualizations of genes **(B)** in cluster 1 related to cell morphogenesis, cell growth, and epithelial differentiation, and in cluster 5 related to immune-related pathways, such as lymphocyte activation, innate and adaptive immune responses, NF-κB signaling, and others. **C.** Gene Set Enrichment Analysis (GSEA) plots for hallmark gene sets of the biological pathways enriched in DES exposure groups compared to the controls across all developmental stages.

A complementary Gene Set Enrichment Analysis (GSEA) across developmental stages confirmed that 1 µg/kg/day of DES exposure perturbs the estrogen signatures in PND46 and PND90 and the androgen response at PND21 (Fig. 6C). Similarly, interferon- *α* / *γ* and IL6–JAK–STAT3 immune pathways were also dysregulated at all developmental stages, but in opposite ways between PND21 and PND90, compared with PND46, suggesting an early immune-hormonal activation, followed by a deactivation and a reactivation (Fig. 6C). In opposite, the dominant enriched pathways at PND46, associated with fatty acid metabolism, adipogenesis, cell cycle regulation, and epithelial-mesenchymal transition, were down-regulated primarily at the two other developmental stages (Fig. 6C). Interestingly, this up-regulation in pathways associated with adipogenesis and metabolism directly correlates with the observed increases in adipocyte number and altered fatty acid concentration aforementioned (Fig. 2–3). However, the development of the mammary gland may be impacted by the enrichment in pathways linked to the innate and adaptive immune system through cytokines and chemokines. Together, these results demonstrate that exposure to 1 μg/kg/day of DES *in utero* triggers a developmental stage-dependent transcriptional change of the mammary gland, initiating with hormonal and immune activation, shifting to metabolic and adipogenic reprogramming at peri-puberty, and maintaining enhanced epithelial growth signatures into adulthood.

### Temporal remodeling of the mammary gland immune microenvironment following in utero DES exposure across developmental stages

The association of cluster 5 gene signatures with immune activation and the enrichment of immune-related pathways at PND21 and PND90 (Fig. 6A-C) led us to investigate the immune profiles of the mammary glands. To do so, we applied the GSEA M8 cell-type signature gene analysis, which predicted strong enrichment of T-cell, B-cell, macrophage, monocyte, and dendritic cell signatures in DES-exposed glands at PND21 and PND90 (Fig. 7A). In contrast, the DES-exposed group at PND46 showed predominant enrichment of stromal fibroblast, basal/myofibroblast, endothelial, and epithelial cell signatures, in agreement with our earlier stromal tissue analyses (Fig. 7A).

**Figure 7.**
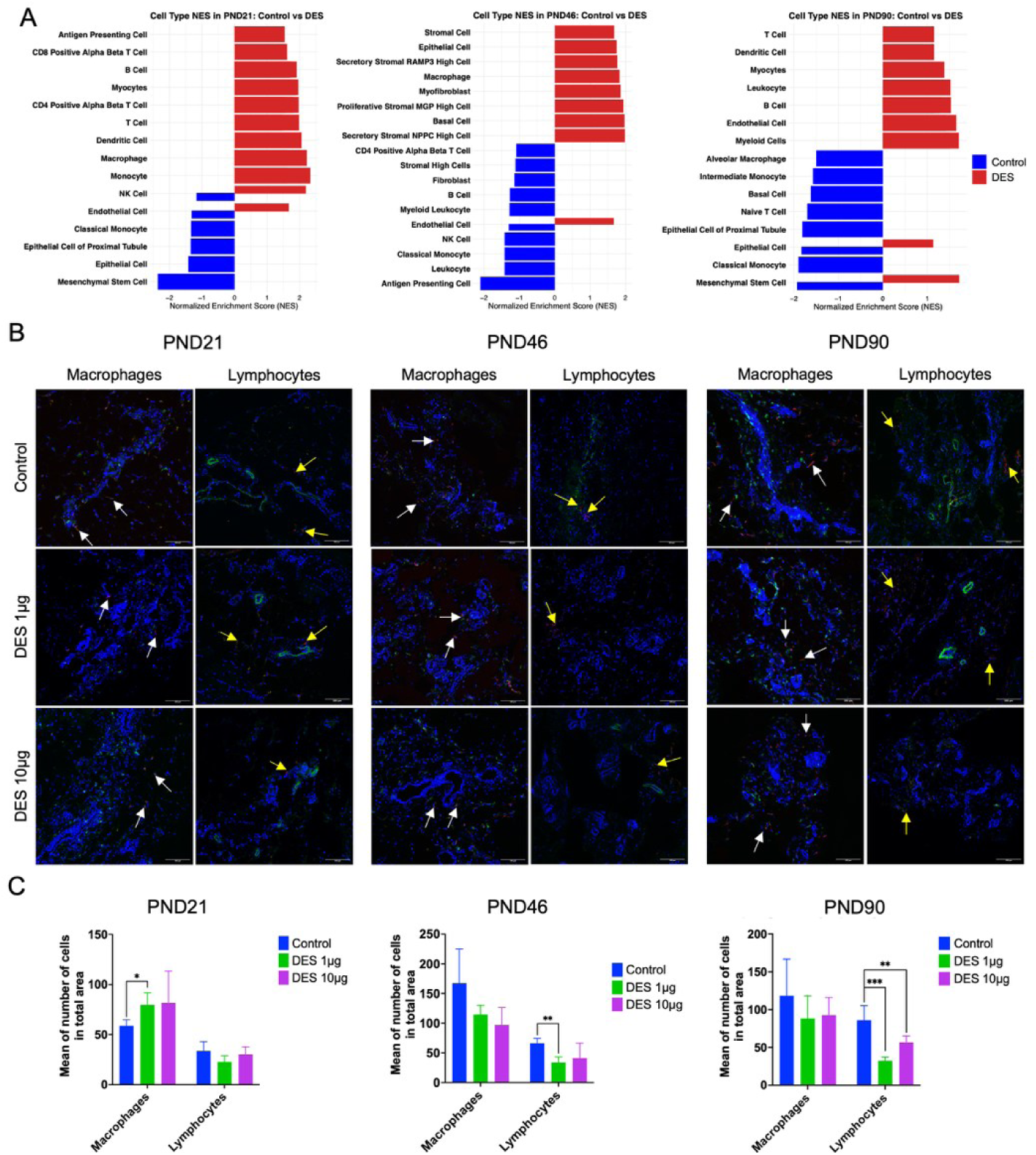
*In utero* DES exposure (1 and 10 ug/kg/day) alters the cell type enrichment and immune cell quantification in mammary glands. **A.** GSEA analysis using the M8 cell type gene sets identified cell types enriched in DES-treated groups (1 ug/kg/day) compared to controls across developmental stages. **B.** Immune cell quantification by immunofluorescence staining using markers of macrophages (CD45 (green) and CD68 (red)), and of lymphocytes (CD45 (green) and CD3 (red)). Magnification 10x, scale bar 300 μm. White and yellow arrows point to macrophages and lymphocytes, respectively, n=5 per group. **C.** Histogram representation of macrophage and lymphocyte number in the total area. Graphs represent the mean of the group with SD (*p < 0.05, **p < 0.01, and ***p < 0.001).

We then validated these *in silico* predictions by immunofluorescent analysis of macrophages and lymphocytes in tissue sections. Macrophage number was significantly elevated at PND21 in the 1 µg/kg/day of DES group compared to the controls, whereas they were slightly, but not significantly, decreased at PND46 and PND90 (Fig. 7C). Macrophages have been associated with a dual role for epithelial cells growth and regression, and the absence of macrophages results in both reduced epithelial cells proliferation (Chua et al. 2010; Stewart et al. 2019). On the contrary, lymphocyte number was not affected at PND21 but significantly reduced at PND46 for the lowest-dose group and remained decreased at both doses at PND90 (Fig. 7B-C and Fig. S7-9). Impacting the cellular, humoral and immunoregulatory branched formation depending on hormones and cytokine signals and the cell-cell interaction (Need et al. 2014). In summary, these data demonstrate that *in utero* DES exposure induces a dynamic stage-specific reprogramming of the mammary gland immune landscape marked by early recruitment of both innate and adaptive immune cells, a peri-puberty shift toward stromal and epithelial cell predominance, and late-stage attenuation of immune infiltration.

### Developmental stage-specific cytokine and chemokine levels modifications in the mammary gland following in utero DES exposure

Given the observed alterations in immune cell populations, we observed changes in macrophages and lymphocytes distribution, and then we next investigated whether *in utero* DES exposure also perturbed the cytokine and chemokine signaling landscape. On one hand, we investigated the profile of pro-inflammatory cytokines as G-CSF, GM-CSF, INF-γ, IL-1α, IL-1β, IL-2, IL-5, IL-6, IL-12p70, IL-17A, and TNF, some anti-inflammatory cytokines as IL-4, IL-10, and IL-13. On the other hand, pro-inflammatory chemokines as CCL1/Eotaxin, CXCL1/GRO-α, CXCL10/IP-10, CCL2/MCP-1, CCL7/MCP-3, CCL3/MIP-1α, CXCL2/MIP2, and CCL5/RANTES. These could be produced by epithelial cells, adipocytes, fibroblasts, and immune cells (Need et al. 2014; Stewart et al. 2019; Wilson et al. 2020). To this end, we quantified 14 cytokines and 8 chemokines using a multiplex Luminex assay across all developmental stages.

PCA revealed distinct developmental stages and dose-specific clustering patterns. At PND21, cytokine vectors predominantly aligned with the control group, whereas chemokine vectors were more closely associated with the DES 1 µg/kg/day group driving the remodeling of immune cells recruitment in response to the low dose (Fig. 8A). At PND46, a transition in immune signaling was evident, as cytokines clustered directionally with the DES 10 µg/kg/day group. In contrast, chemokines were more aligned with the control (Fig. 8B). By PND90, a robust immune response activation was evident in both DES groups. PCA revealed that both cytokines and chemokines showed a pronounced shift toward the DES 10 µg/kg/day group, indicating enhanced and coordinated immune signaling in adult glands following exposure to the highest dose of DES (Fig. 8C). In correlation with our results, DES 1 µg/kg/day had a stronger effect in increasing some fatty acids distribution across mammary gland development, from immune microenvironment DES 10 µg/kg/day has a stronger effect in the modulation of cytokines and chemokines.

**Figure 8.**
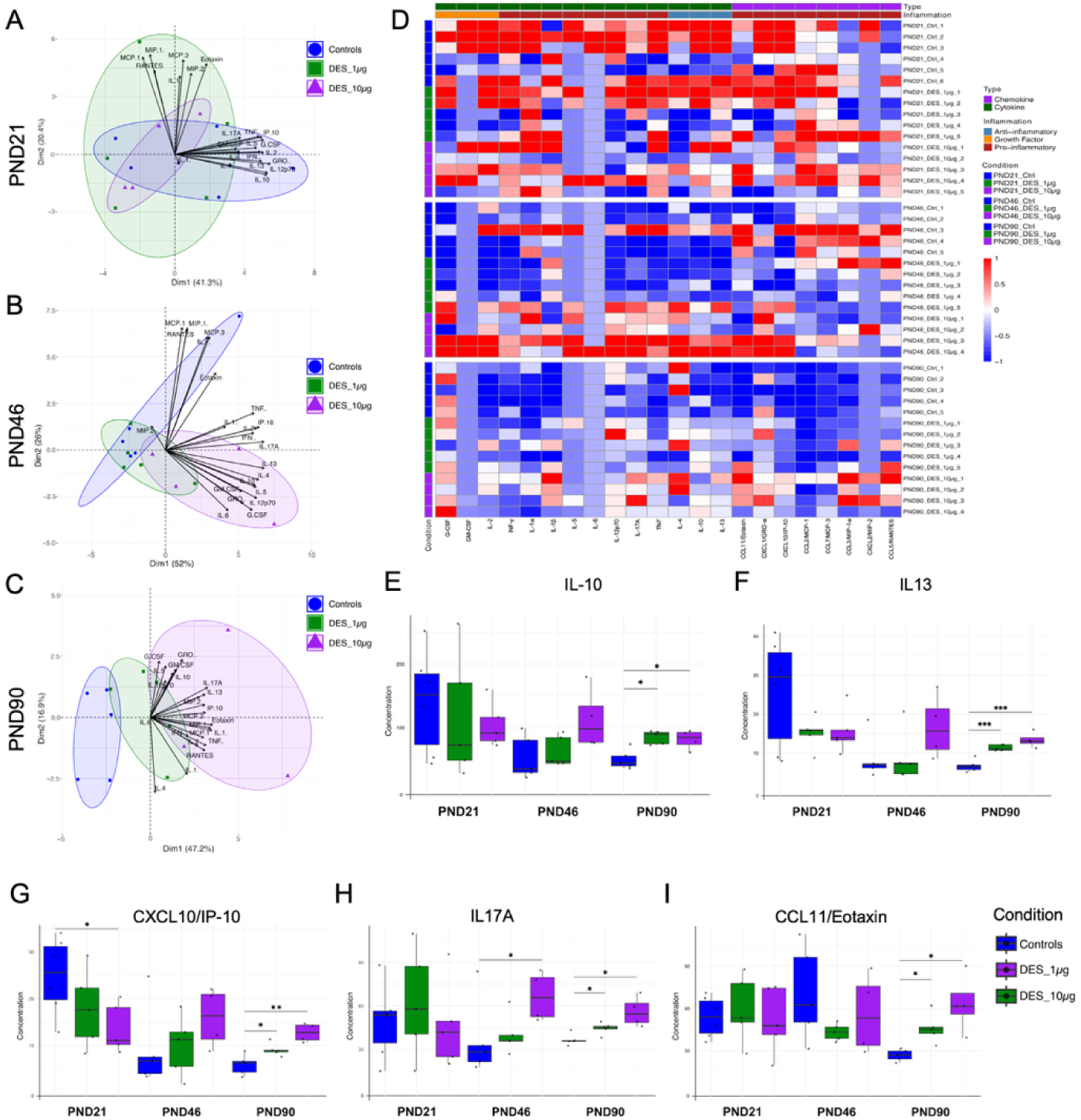
Exposure to DES affects the cytokine and chemokine profiles across different developmental stages. **(A–C)** Principal component analysis of cytokine and chemokine profiles at PND21 (**A**), PND46 (**B**), and PND90 (**C**). (**D**) Heatmap showing relative expression levels of 22 cytokines and chemokines across developmental stages and treatment conditions. (**E–I**) Histograms showing differentially expressed anti-inflammatory cytokines IL-10 (**E**), IL-13 (F), and pro-inflammatory cytokines and chemokines CXCL10/IP10 (**G**), IL-17A (H), and CCL11/Eotaxin (I) in DES-treated groups compared to controls. Data are presented as mean ± SD. *p < 0.05, **p < 0.01, ***p < 0.001.

To further visualize the expression profiles, we generated a heatmap comparing all cytokine and chemokine levels in all groups. At PND21, cytokine expression was higher in control animals compared to DES-exposed groups, particularly for the lowest dose. In contrast, at PND90, cytokine and chemokine expression were markedly elevated in both DES-exposed groups compared to controls (Fig. 8D), supporting a late-stage pro−inflammatory shift induced by *in utero* exposure. Analysis of individual cytokine and chemokine levels confirmed these patterns. At PND21, only CXCL10/IP-10 was significantly downregulated in the DES 10 µg/kg/day group, which could indicate a reduction of CXCR3/GPR9 effector T-cells and NK-cells, and an increase in macrophages numbers at this developmental stage (Fig. 7C). At PND46, GM-CSF, IL-12p70, and CXCL1/GRO-α were upregulated in the highest-dose group, suggesting activation of myeloid and granulocytic programs for monocytes/macrophages, neutrophils or eosinophils (Fig. 8 and Table 2). On the contrary, CCL2/MCP-1 was downregulated, which could indicate a limited monocyte infiltration, a neutrophil recruitment, and/or resident macrophage persistence (Table 2) (Deshmane et al. 2009).

**Table 2.**
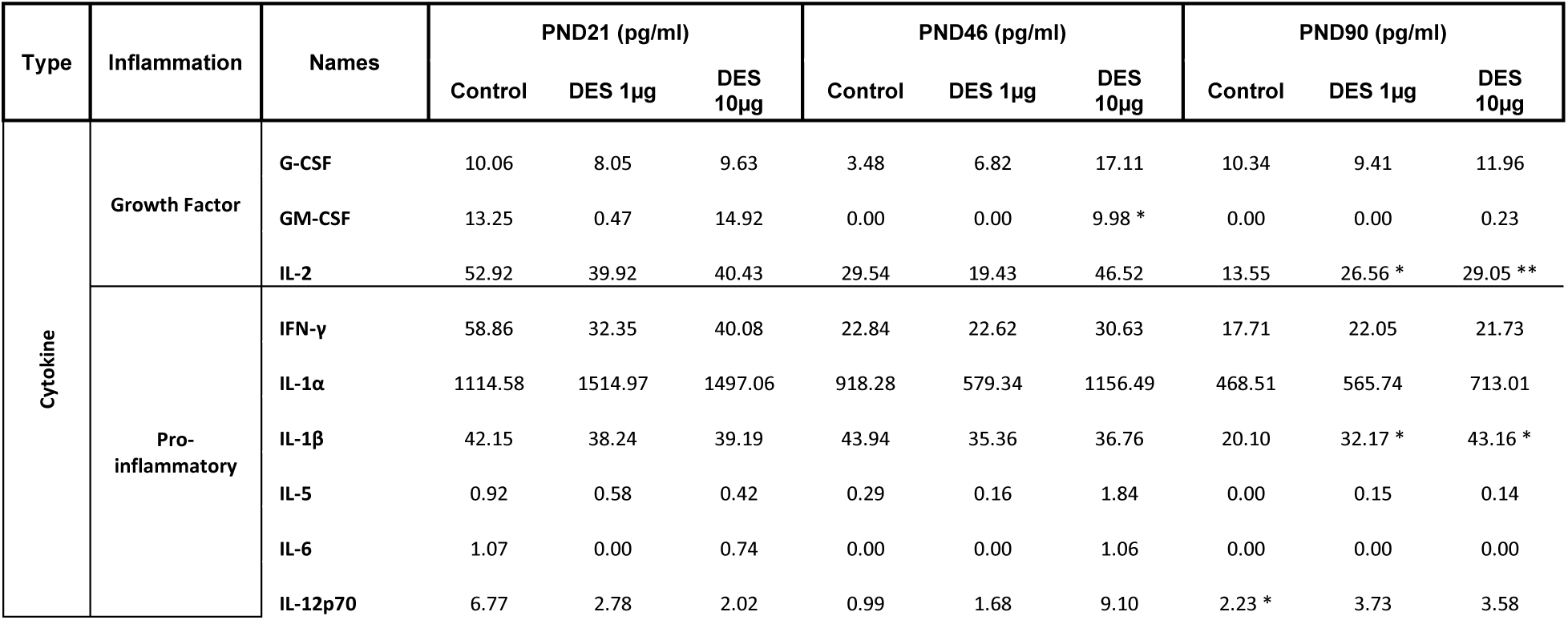
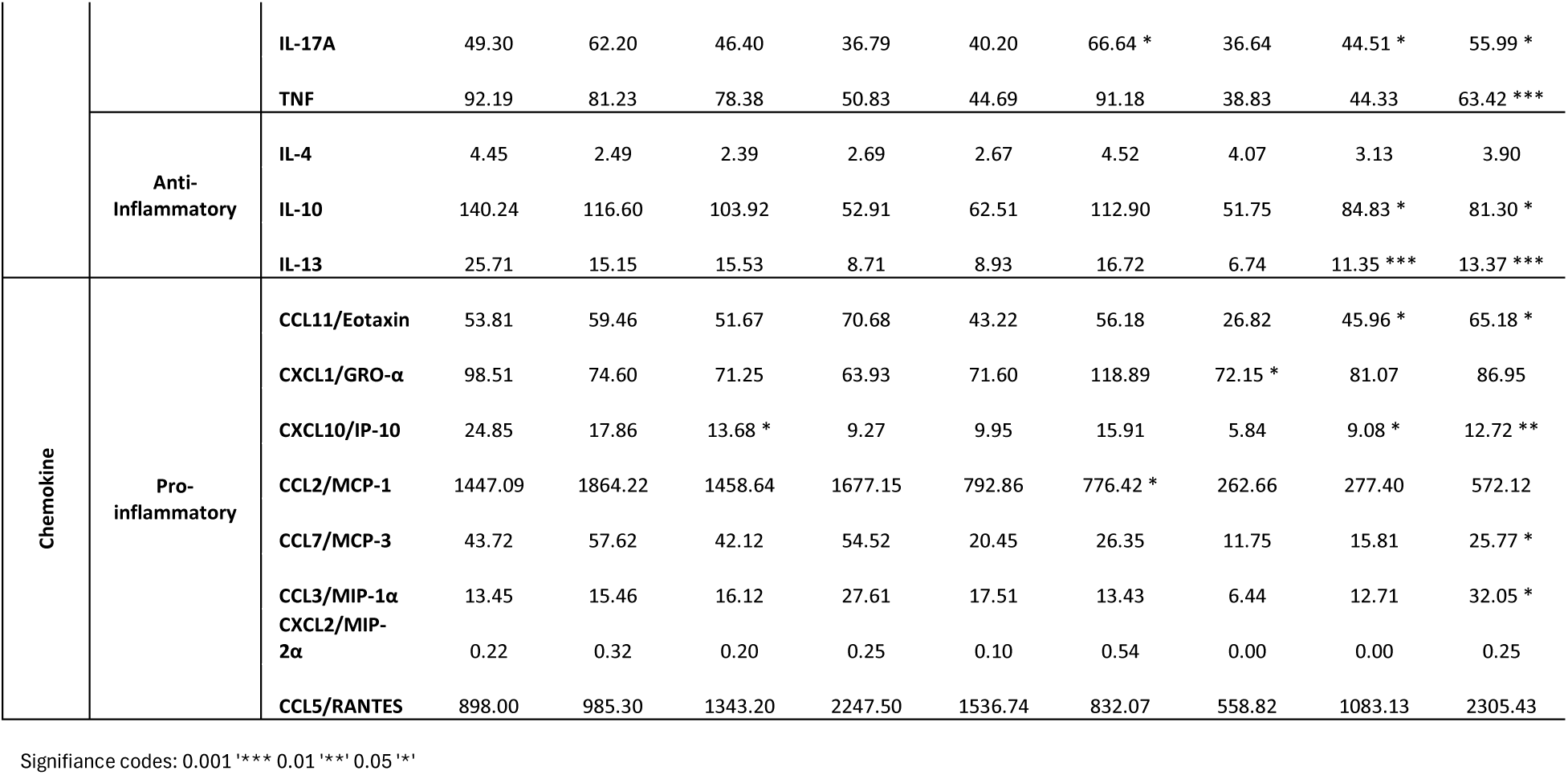
Cytokine and chemokine profiles across developmental stages and DES exposure groups.

In marked contrast, PND90 showed an extensive upregulation of immune mediators in both DES groups. This was characterized by elevated levels of pro-inflammatory cytokines (IL-1β, IL-2, TNF, IL-17A), regulatory/Th2-associated cytokines (IL-10, IL-13), and chemokines involved in broad leukocyte recruitment (CXCL10/IP10, CCL11/Eotaxin, CCL3/MIP-α, CCL7/MCP-3) (Fig. 8E–I). These patterns point to a complex immune microenvironment, marked by persistent low-grade inflammation, altered leukocyte composition, and remodeling-prone signaling in adult DES-exposed glands. Collectively, these results indicate that *in utero* DES exposure induces a temporal remodeling of the mammary gland cytokine and chemokine environment. While early life exposure transiently suppresses select chemokines, late-stage responses are characterized by a robust, dose-dependent activation of both pro-inflammatory and immunoregulatory signals, likely contributing to the altered stromal and epithelial landscape observed at PND90.

## Discussion

### Effects of DES on epithelial development

Estrogens are known to regulate epithelial proliferation and ductal morphogenesis primarily through ER *α* signaling pathways (Kanaya et al. 2019; Rusidze et al. 2021). Mallepell *et al*. demonstrated that ERα knockout (KO) mice showed a decrease in TEB formation and ductal elongation at 4 and 10 postnatal weeks (Mallepell et al. 2006). Conversely, neonatal administration of DES (1 µg) significantly increases TEB number at PND50 (Ninomiya et al. 2007). Umekita *et al*. further demonstrated that a single neonatal dose of DES (0.1 – 100 µg) elevates TEB counts at PND39 and PND49 and upregulates lactation-associated genes (β-casein, γ-casein, and WAP) (Umekita et al. 2011). In contrast, *in utero* and lactational exposure to DES (6 µg) delays mammary differentiation and alters milk production in the offspring (Kass et al. 2012), demonstrating that the time of exposure has an important impact on the consequences later in life (Fenton 2006).

The mammary epithelium begins around GD12 with the formation of placodes that will elongate to form the canals around GD16 (Fenton 2006; Macias and Hinck 2012). ERα has been found in mammary mesenchymal cells surrounding this growing epithelium, likely helping in the formation of the rudimentary primordial epithelial tissue between GD14-16 (Shyamala et al. 2002). For this reason, we chose to expose rats to DES at the end of gestation; this time is also thought to correspond to the period of differentiation of the fetal mammary gland stroma (Líška et al. 2016; Masso-Welch et al. 2000). Although this exposure to DES has little impact on epithelium development, our results suggest that *in utero* exposure to DES at 10 µg/kg/day reduced the complexity of ductal architecture in adult rats. Interestingly, a recent study from Montévil *et al*., reported that rats’ exposure to 6 and 12 µg of DES during gestation and through lactation also reduces the area of epithelial tissue and the complexity at PND22 (Montevil et al. 2025), but adult rats were not assessed in that study. However, it suggests that effects are more significant when the exposure spans a longer period during perinatal life. Notably, in our study, the stroma compartment exhibited even more pronounced alterations than the epithelium, and the effects were generally more critical at the lowest dose. Our findings suggest that gestational exposure to DES disrupts the epithelial and stromal tissue, leading to long-term consequences that persist into adulthood.

### Developmental stage disruption of stromal adipogenesis and lipid remodeling upon in utero exposure to DES

One of the most common results was an increase in the number of adipocytes in groups exposed to DES, and a slight increase in their size at PND90 for the lowest dose. Concurrently, exposure to the lowest dose of DES deregulated 10 of 43 fatty acids across development, elevating medium and long chain, saturated and key PUFAs at PND46 and PND90, whereas exposure to the highest dose selectively reduced lauric and myristic acids at PND21, indicating that DES may cause stromal changes through modified lipid production and storage. Indeed, lauric acid has been shown to activate PI3K/Akt/mTOR signaling, promoting the proliferation of mouse mammary epithelial cells and alveolar development during lactation (Meng et al. 2017; Yang et al. 2020). Moreover, elevated docosapentaenoic acid has been linked to lobulo-alveolar differentiation and upregulation of β-casein in murine models, reflecting a premature, pregnancy-like stromal phenotype in the epithelial tissue (Liu et al. 2007). Together, these data indicate that DES low-dose exposure induces persistent changes of both saturated and unsaturated fatty acids during peri-puberty and adulthood.

Interestingly, results observed upon *in utero* DES exposure mimic, in part, results that we obtained previously when we exposed rats to di(2-ethylhexyl)phthalate (DEHP) or 1,2-cyclohexane dicarboxylic acid diisononyl ester (DINCH) during gestation and lactation (Crobeddu et al. 2022). Indeed, DEHP and DINCH exposure had little effect on the epithelium but resulted in decreased adipocyte size, as well as reduced triglycerides and free fatty acid levels at PND90 (Crobeddu et al. 2022). In addition, as for DES, opposite effects were observed between PND46 and PND90. To our knowledge, only one study has examined the effects of DES on mammary gland stroma in parallel to bisphenol A (BPA). In mice exposed to a high dose of DES (100 µg) from GD9 to GD18, an increase in gland stiffness was reported in adult mice (Wormsbaecher et al. 2020). Interestingly, *in utero* BPA exposure resulted in fibroblast reprogramming, higher levels of collagens, and increased fiber width (Wormsbaecher et al. 2020). Similarly, Wadia *et al*. reported that in the mammary gland of mice exposed to a low dose of BSA from GD8 to GD19, key stromal and adipogenic genes (*Fabp4* and *PPARγ*) were downregulated later in life (PND18) (Wadia et al. 2013). These results, when combined, demonstrate the sensitivity of mammary stromal development to early life exposure to DES and other endocrine-disrupting chemicals. They also underscore the long-term impact of *in utero* exposure on the structure and function of the mammary gland.

### Developmental stage-specific disruption of stromal organization

Another main component of the stroma that fluctuates during mammary gland development and actively participates in epithelium development is the ECM. Brownfield *et al*. demonstrated that precise collagen I fiber orientation is essential for ductal branching morphogenesis via Rac1–MMP2 mediated matrix remodeling in 3D cultures (Brownfield et al. 2013). In addition, previous studies showed that laminin-rich matrices drive epithelial polarization and acinar lumen formation (Gudjonsson et al. 2002). Although we did not find any difference when looking at proportions of collagen versus epithelial and adipocyte components, immunofluorescence staining revealed an early depletion of collagens that is significant for I/V at PND21 (1 µg), followed by a transient increase in collagen I and laminin at PND46 (10 µg), and return to control levels at PND90. To further understand these contradictory results, we used SHG imaging to evaluate the number and orientation of collagen fibers. First, when looking only at control animals, we observed a developmental increase in fiber number between PND21 and PND46, followed by a stabilization, which is consistent with ductal elongation (Buchmann et al. 2021). We also observed a modification in the orientation as reflected by a predominance in fibers with an angle between 30 and 90 degrees at PND46 and PND90. Brownfield *et al*. found that collagen I orientation modulated dramatic epithelial co-orientation encoded architectural branch orientation (Brownfield et al. 2013). Nevertheless, overall, DES seems to induce a dynamic, developmental stage-dependent ECM reorganization that could explain the decrease in epithelium complexity observed at PND90. More experiments are needed to understand the links between fiber orientation and mammary gland development.

### Developmental stage-specific impact on the immune microenvironment

The pronounced stromal remodeling prompted us to investigate accompanying changes in the mammary immune landscape. Our GSEA M8 cell-type analysis predicted early enrichment of immune cells (T-cells, B-cells, macrophages, monocytes, and dendritic cells) at PND21 and PND90, followed by stromal and epithelial abundance at PND46 in the exposure group. These *in silico* predictions align with single-cell findings by Kanaya *et al*., who reported that estrogenic stimulation in adult mice expands immune clusters, including T/NK-cells, antigen-presenting cells (APC), and M2-like macrophages (Kanaya et al. 2019).

Immune cells are known to play roles in tissue remodeling, wound healing, and the clearance of apoptotic cells (Need et al. 2014; Reed and Schwertfeger 2010). Macrophages have been associated with compiling collagen into long fibers around the TEBs during the ductal elongation (Reed and Schwertfeger 2010). Eosinophils have been linked with facilitating bifurcation during TEBs formation (Gouon-EvansRothenberg and Pollard 2000; Reed and Schwertfeger 2010). Lymphocytes CD4+ and CD8+ have been found adjacent to APC in the mammary ducts (Plaks et al. 2015). Consistent with this, our immunofluorescence data confirmed that DES induced alterations in immune cell populations. Notably, DES-exposed glands showed a significant increase in CD68⁺ macrophage density at PND21, but a marked decrease in CD3⁺ T-lymphocyte density at PND46 and PND90 compared to controls. These changes suggest that DES perturbs the temporal recruitment and retention of immune populations, possibly creating a chronic low-grade inflammatory environment that can influence ECM remodeling, potentially altering phagocytic clearance of apoptotic cells and remodeling signals during ductal morphogenesis (Hitchcock et al. 2022; Need et al. 2014).

This immunological reprogramming is strongly coupled to stromal remodeling, adipocyte expansion, and epithelial changes, highlighting the immune microenvironment as a critical mediator of estrogenic disruption. During PND21, DES exposure was associated with reduced expression of CXCL10/IP-10, associated with T-cell and NK-cell chemoattractant, despite elevated macrophage density (Hirani et al. 2023). This change in cytokines could indicate that macrophage infiltration in the absence of T-cell and NK-cell recruitment may result in macrophage polarization, favoring tissue-remodeling (M2-like macrophages) rather than immune-regulatory phenotypes (Reed and Schwertfeger 2010). Previous studies demonstrated in CSF-1 null mutant mice that the absence of macrophages in terminal end buds altered their forms and collagen degradation (Ingman et al. 2006; Lin et al. 2002). However, at PND46, the DES-induced immune landscape shifted toward a myeloid-dominant activation profile, as evidenced by upregulation of GM-CSF, IL-12p70, and CXCL1/GRO-α. These cytokines are known to support dendritic cell maturation, neutrophil recruitment, and stromal remodeling (Shiomi and Usui 2015). This altered immune environment coincides with the peak in adipogenesis and matrix protein expression, reinforcing the concept that the immune system acts as a modulator of stromal composition and organization during pubertal morphogenesis (WynnChawla and Pollard 2013). Stein *et al*. showed the presence of CXCL1/GRO-α on day 1 of the involution process, followed by genes associated with macrophage chemoattraction and differentiation on days 3 and 4 (Stein et al. 2004). These changes suggest that DES disrupts the immune cell signaling, redirecting immune-stromal crosstalk in a way that may reshape the mammary gland during pre-puberty and peri-puberty.

At PND90, DES-exposed mammary glands exhibited a complex cytokine signature that included IL-1β, IL-17A, TNF, IL-10, and IL-13, alongside potent chemokines like CCL3/MIP-1α, CCL7/MCP-3, and CXCL10/IP-10. This profile reflects a chronic, unresolved immune activation with features of Th1, Th2, and Th17 polarization. The elevated IL-17A concentration, in particular, is indicative of tissue inflammation and epithelial stress, and has been linked to fibrosis and aberrant ECM turnover in systemic sclerosis (Dufour et al. 2020). Th1 cells, located near the growth of mammary ducts, secrete pro−inflammatory cytokines such as TNF and IFN-γ, which can negatively influence ductal growth, while Th2 cells secrete cytokines like IL-4 and IL-10, associated with tissue remodeling (Sakaguchi et al. 2008). Fischer *et al*. demonstrated that *in utero* exposure to BPA alters mammary cytokine profiles, significantly downregulating several chemokine genes (*Cxcl2, Cxcl14, Ccl20*), interleukins *Il1β*, and *Il7r receptor*, and interferon-pathway genes in adult mammary tissue (Fischer et al. 2016). Such dysregulation of both pro- and anti-inflammatory cytokines can create a microenvironment that favors persistent inflammation while suppressing effective immune surveillance. In our DES exposure group, the cytokine perturbations at PND90 further support the notion of a sustained immunomodulatory state in the mammary stroma. This altered state, characterized by macrophage enrichment, T-cell paucity, and cytokine imbalance, may facilitate abnormal ECM deposition, orientation, and remodeling, potentially compounding the effects of DES on gland development.

## Conclusion

Overall, our results demonstrate that *in utero* exposure to DES induces developmental stage and dose-specific alterations in mammary gland composition, particularly affecting the stromal compartment. We observed changes in adipocyte number and lipid profiles, transcriptional shifts in adipogenesis and immune signaling pathways, and dynamic remodeling of the ECM. These effects were accompanied by stage-specific disruptions in immune cell populations and cytokine expression, with a pronounced inflammatory and stromal remodeling profile at adulthood. Overall, this study highlights that DES exposure during late gestation persistently reprograms mammary gland development, establishing long-lasting structural and immunological alterations that may impact tissue homeostasis.

## Acknowledgments

Eva Gierski is thanked for their technical assistance.

---

**Supplementary Table 1.**
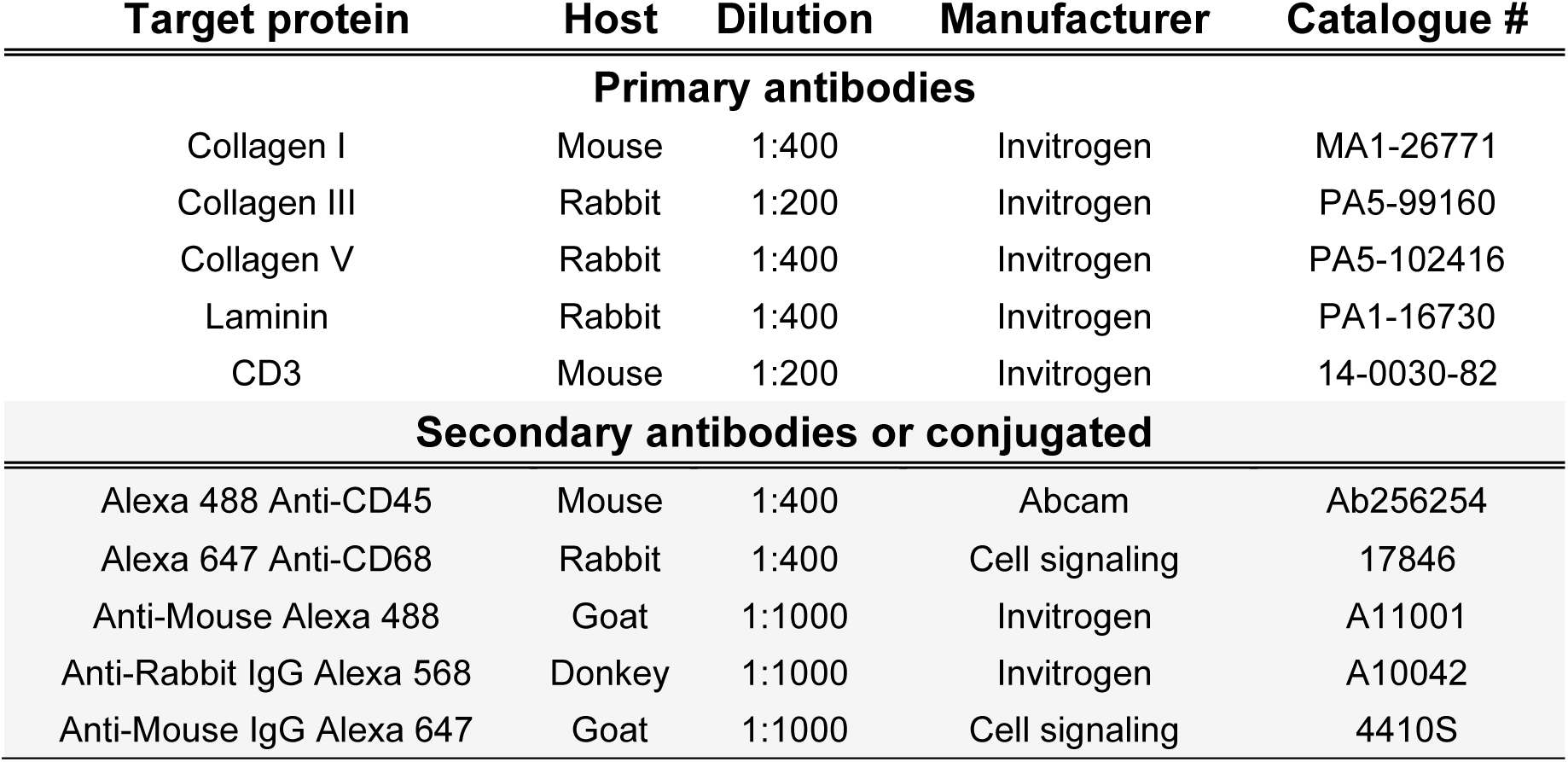
List of Antibodies used for immunofluorescence.

| Target protein | Host | Dilution | Manufacturer | Catalogue # |
| --- | --- | --- | --- | --- |
| **Primary antibodies** |  |  |  |  |
| Collagen I | Mouse | 1:400 | Invitrogen | MA1-26771 |
| Collagen III | Rabbit | 1:200 | Invitrogen | PA5-99160 |
| Collagen V | Rabbit | 1:400 | Invitrogen | PA5-102416 |
| Laminin | Rabbit | 1:400 | Invitrogen | PA1-16730 |
| CD3 | Mouse | 1:200 | Invitrogen | 14-0030-82 |
| **Secondary antibodies or conjugated** |  |  |  |  |
| Alexa 488 Anti-CD45 | Mouse | 1:400 | Abcam | Ab256254 |
| Alexa 647 Anti-CD68 | Rabbit | 1:400 | Cell signaling | 17846 |
| Anti-Mouse Alexa 488 | Goat | 1:1000 | Invitrogen | A11001 |
| Anti-Rabbit IgG Alexa 568 | Donkey | 1:1000 | Invitrogen | A10042 |
| Anti-Mouse IgG Alexa 647 | Goat | 1:1000 | Cell signaling | 4410S |

**Figure Supplementary 1.**
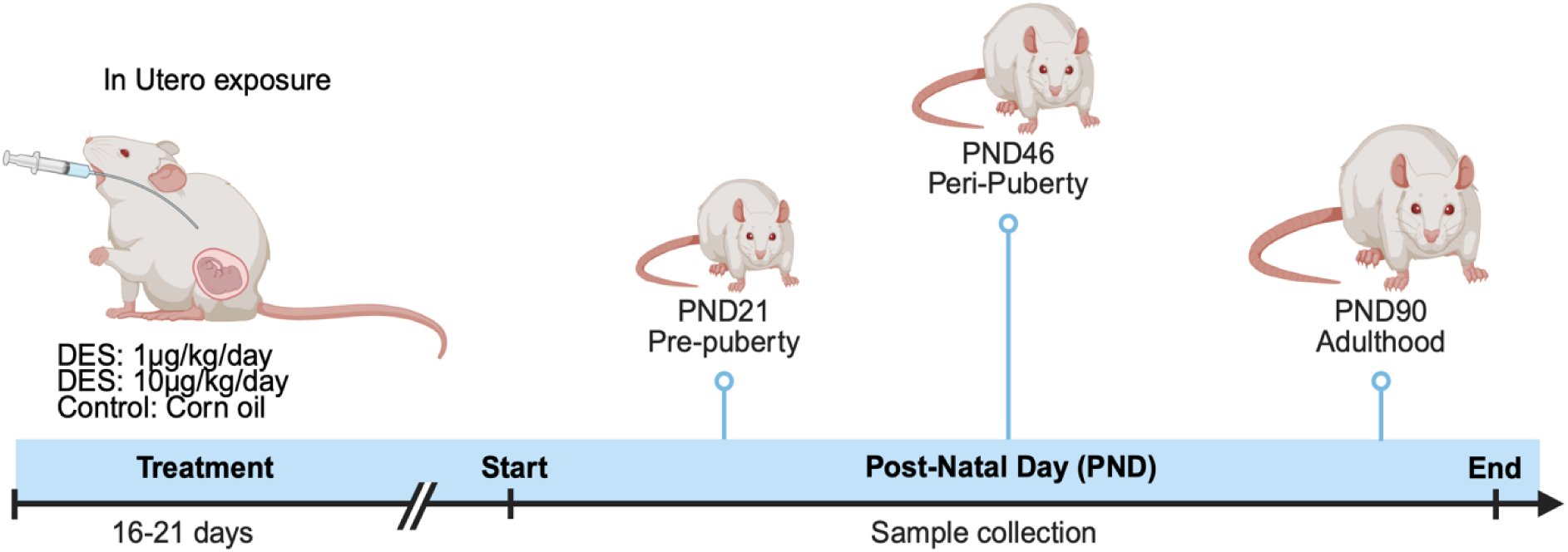
Schematic timeline of *in utero* exposure. Rats were exposed to DES during gestation, and mammary gland samples were collected at pre-puberty (PND21), peri-puberty (PND46), and adulthood (PND90).

**Figure Supplementary 2.**
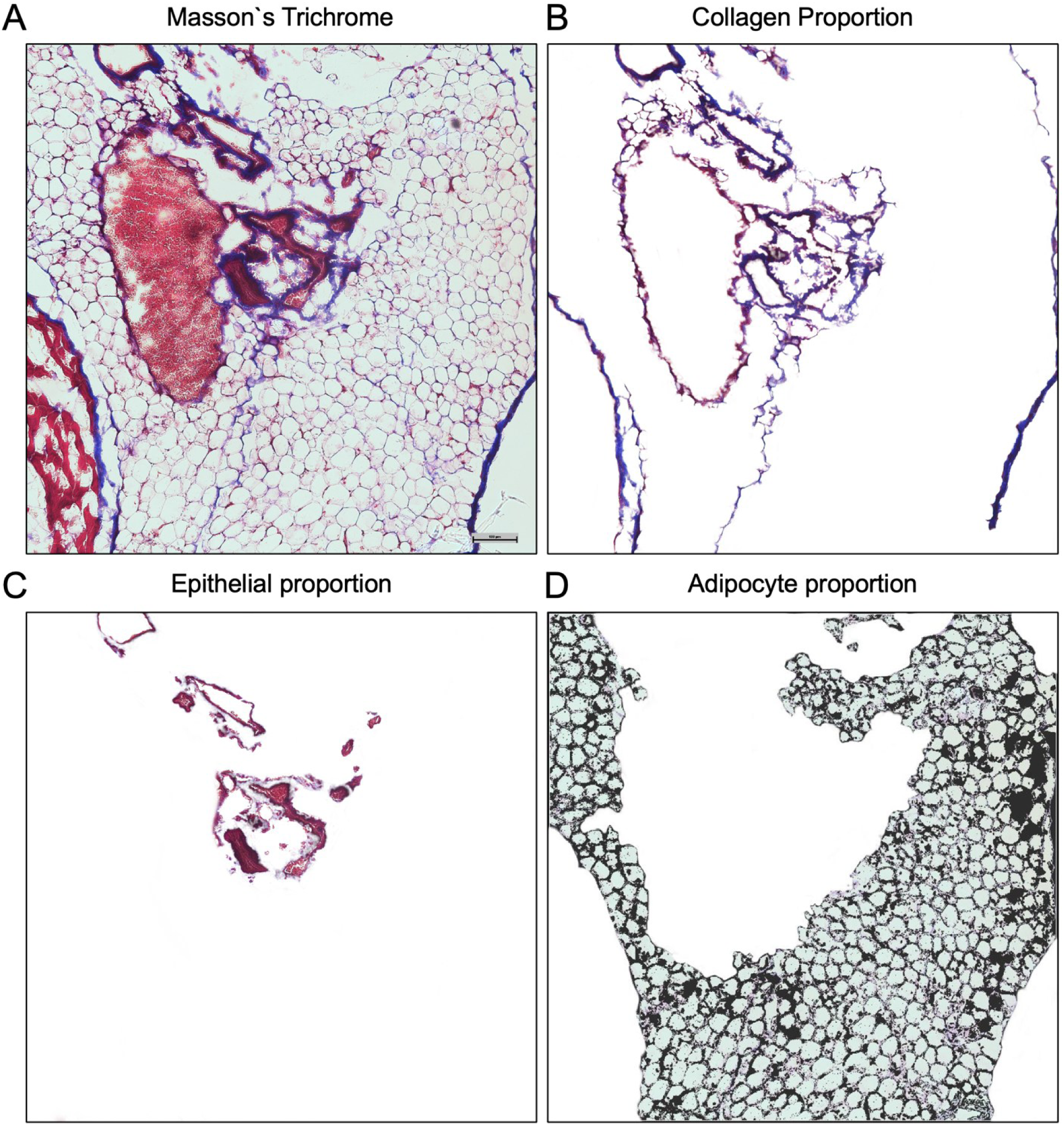
Representative pixel segmentation analysis of a control animal (PND21) by hue, saturation, and intensity.

**Figure Supplementary 3.**
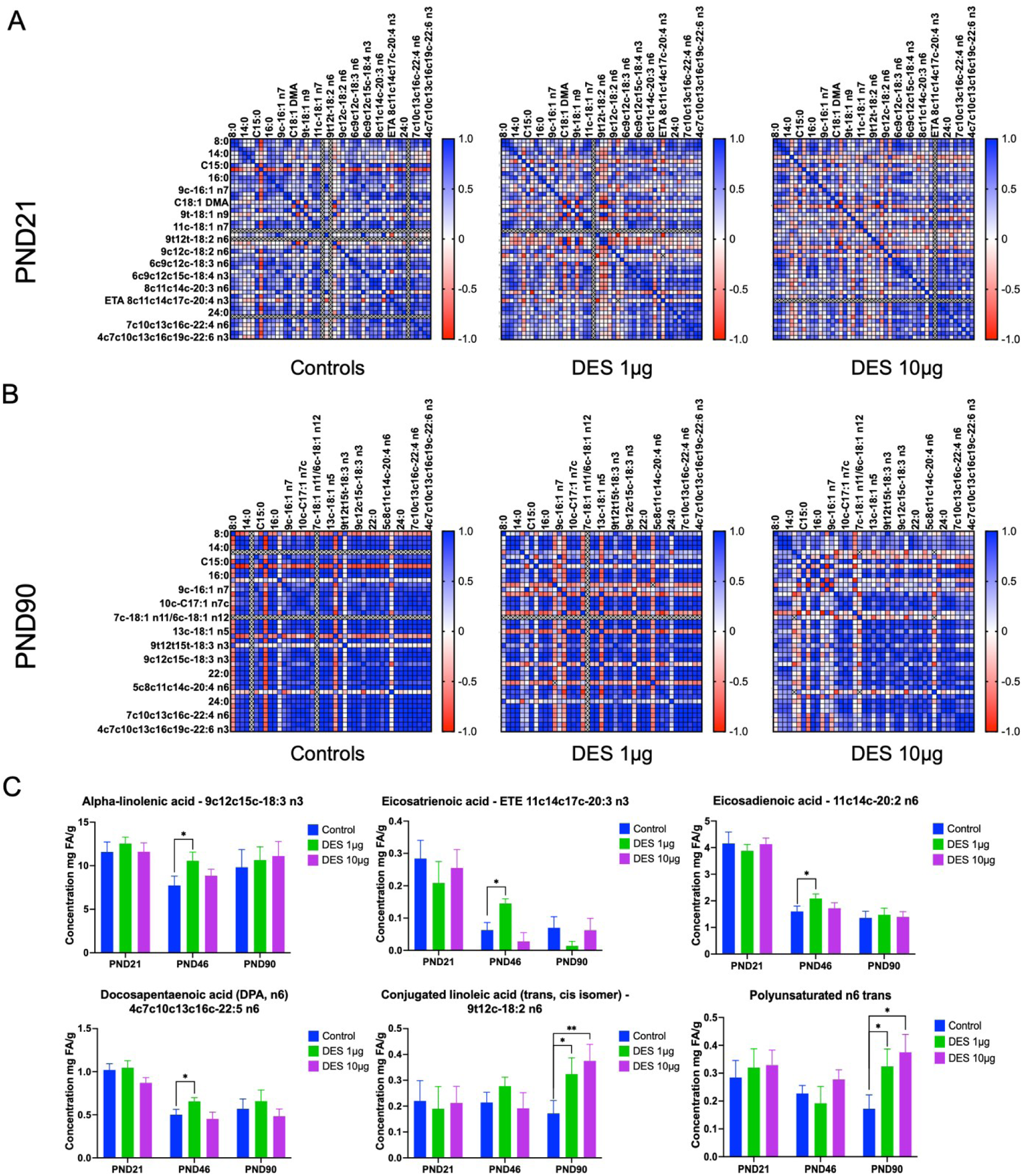
Fatty acid distribution after DES exposure in the mammary gland across developmental stages. A-B. Correlation of fatty acids showing the different lipid profiles at PND46 and PND90 groups after DES exposure. C. Histogram representation of fatty acids. Graphs represent the mean of the group with SD (*p < 0.05).

**Figure Supplementary 4.**
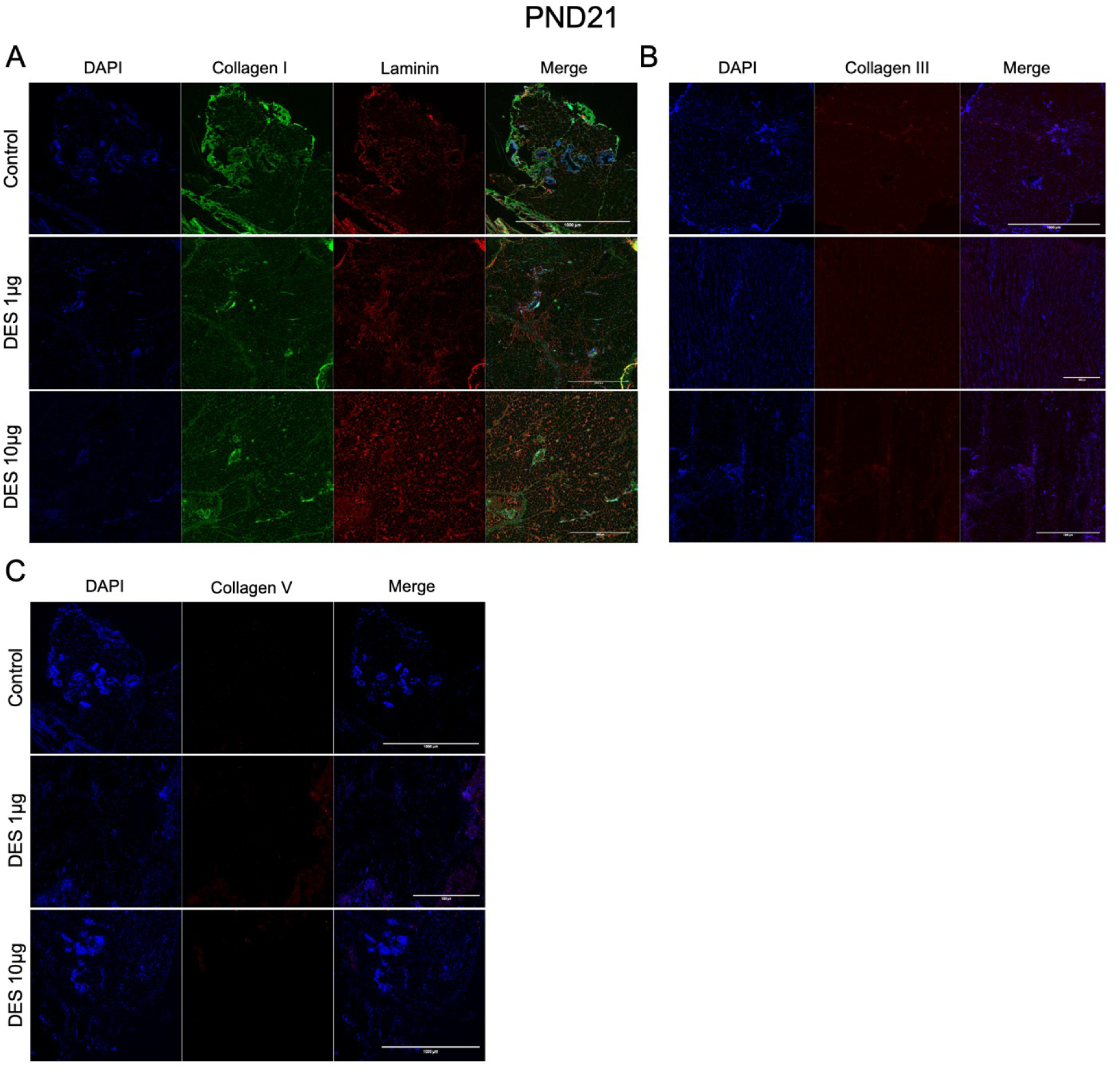
Extracellular matrix remodeling in mammary glands after *in utero* DES exposure at PND21. Representative immunofluorescence images showing Collagen I and laminin (**A**), Collagen III (**B**), and Collagen V (**C**) at PND21 in control and DES-treated groups at 10x magnification (scale bar = 1000μm).

**Figure Supplementary 5.**
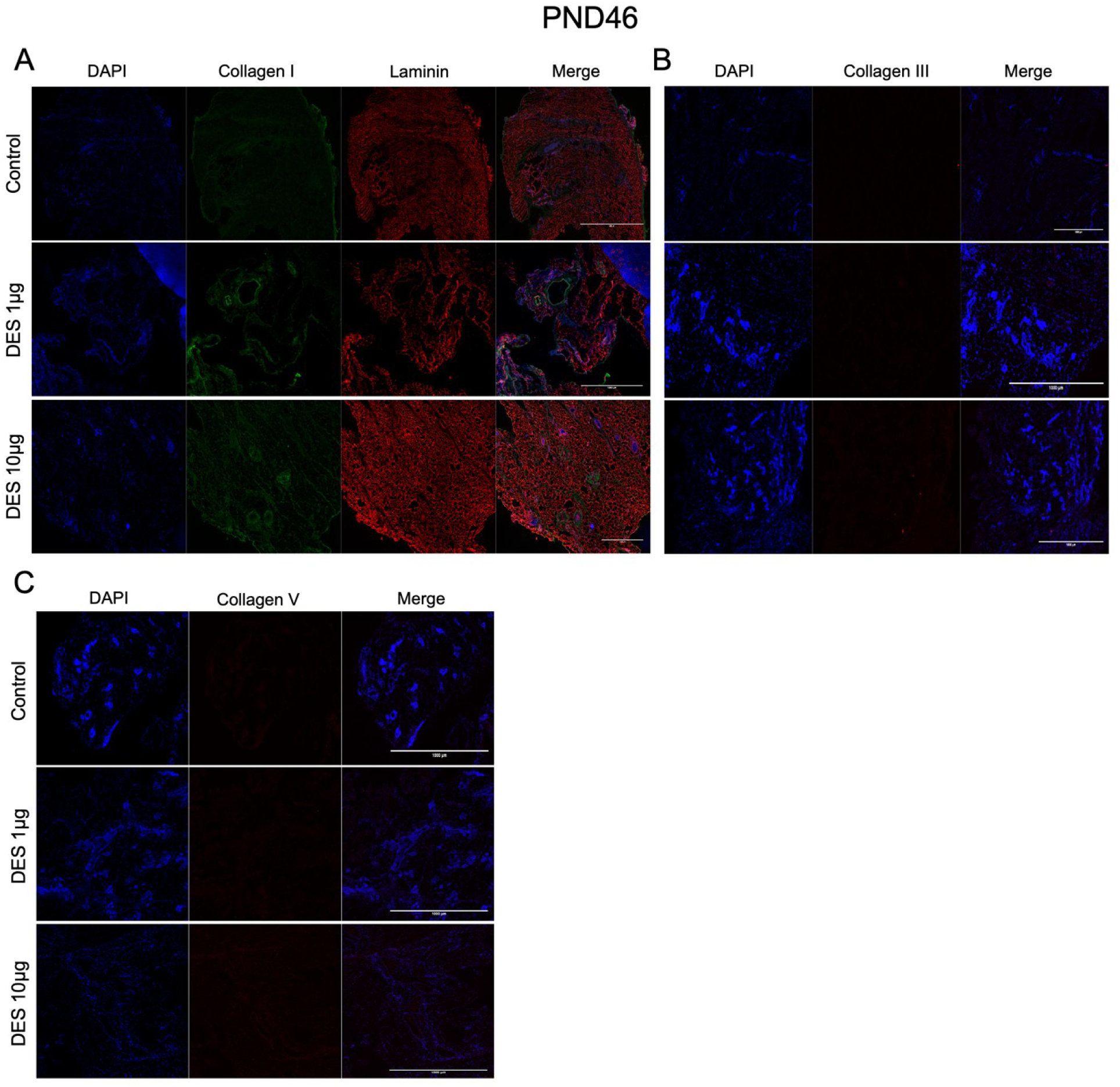
Extracellular matrix remodeling in mammary glands after *in utero* DES exposure at PND46. Representative immunofluorescence images showing Collagen I and laminin (**A**), Collagen III (**B**), and Collagen V (**C**) at PND46 in control and DES-treated groups at 10x magnification (scale bar = 1000μm).

**Figure Supplementary 6.**
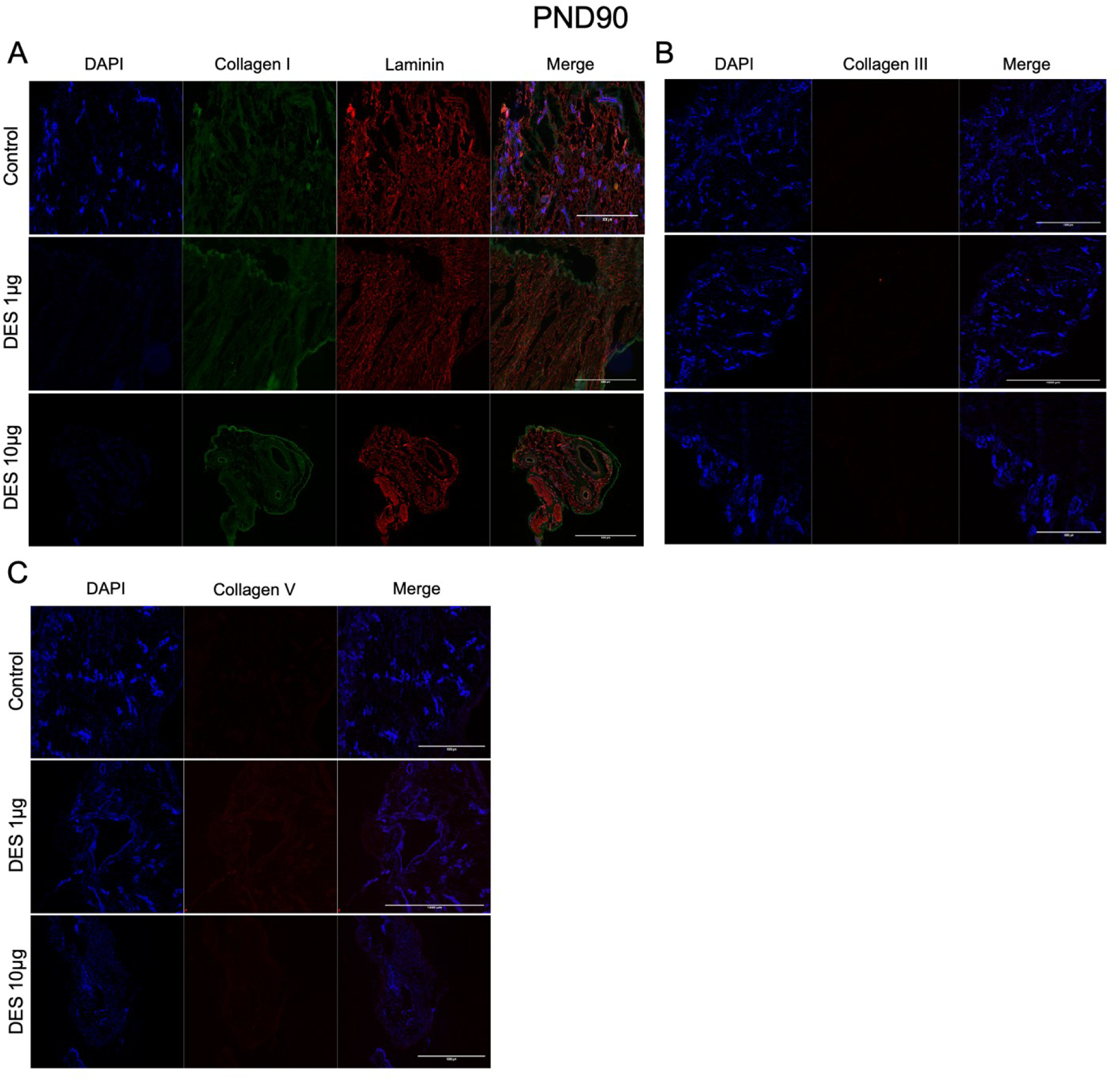
Extracellular matrix remodeling in mammary glands after *in utero* DES exposure at PND90. Representative immunofluorescence images showing Collagen I and laminin (**A**), Collagen III (**B**), and Collagen V (**C**) at PND90 in control and DES-treated groups at 10x magnification (scale bar = 1000μm).

**Figure Supplementary 7.**
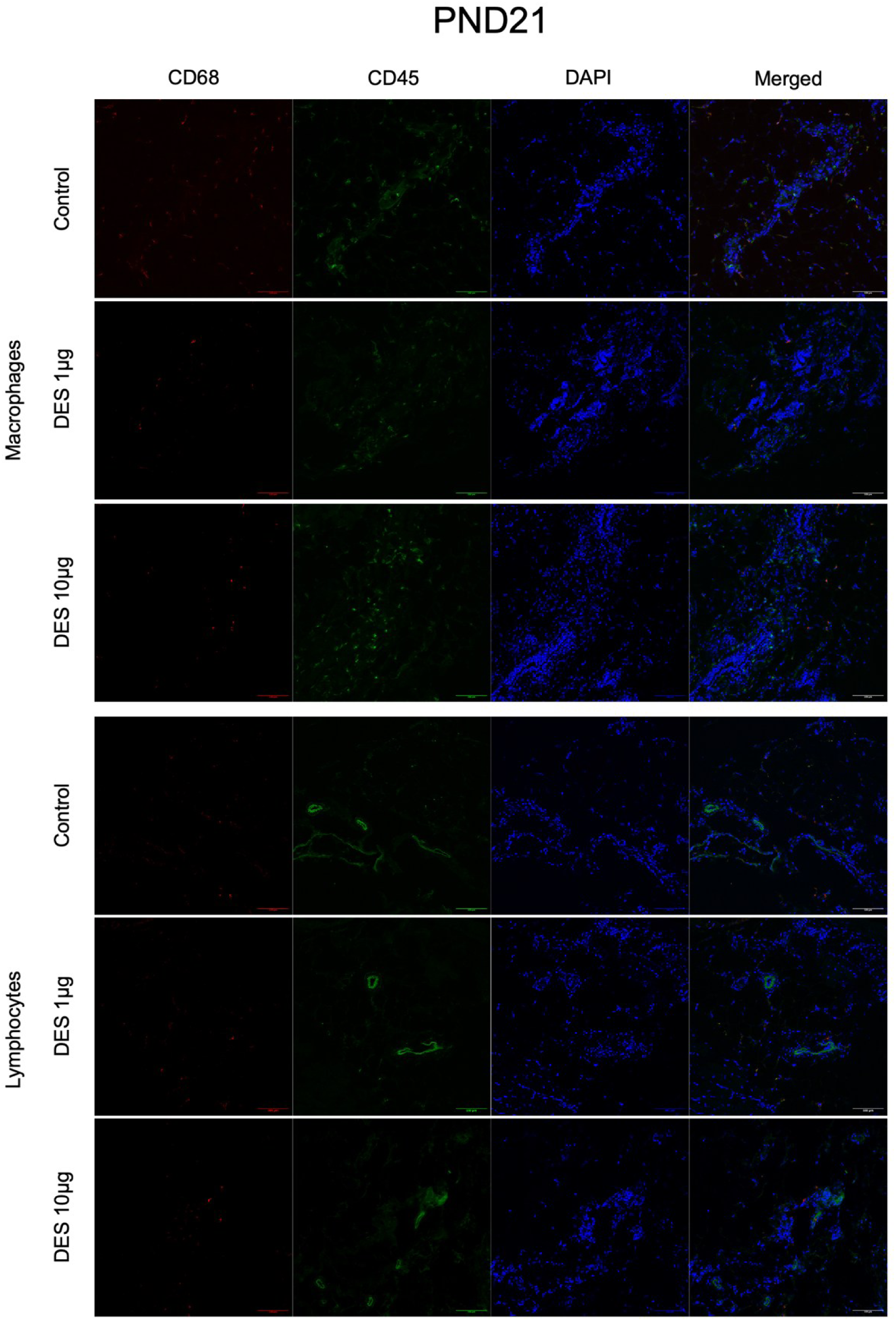
Immune cell quantification at PND21 after DES exposure in the mammary gland. Immunofluorescence staining using markers CD45 and CD68 for macrophages (**A**) and CD45 and CD3 for lymphocytes (**B**). Magnification 10x, scale bar 300 μm.

**Figure Supplementary 8.**
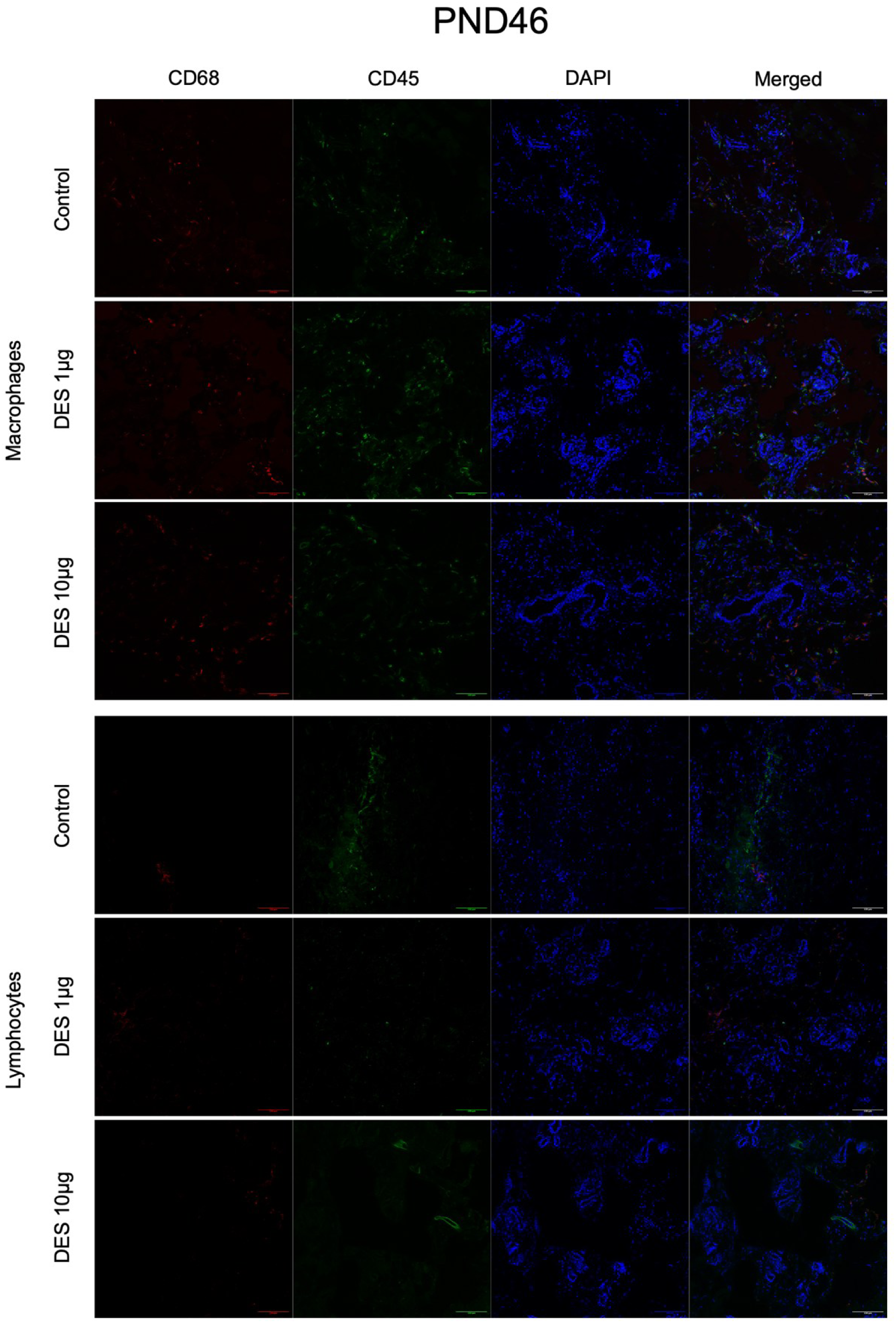
Immune cell quantification at PND46 after DES exposure in the mammary gland. Immunofluorescence staining using markers CD45 and CD68 for macrophages (**A**) and CD45 and CD3 for lymphocytes (**B**). Magnification 10x, scale bar 300 μm.

**Figure Supplementary 9.**
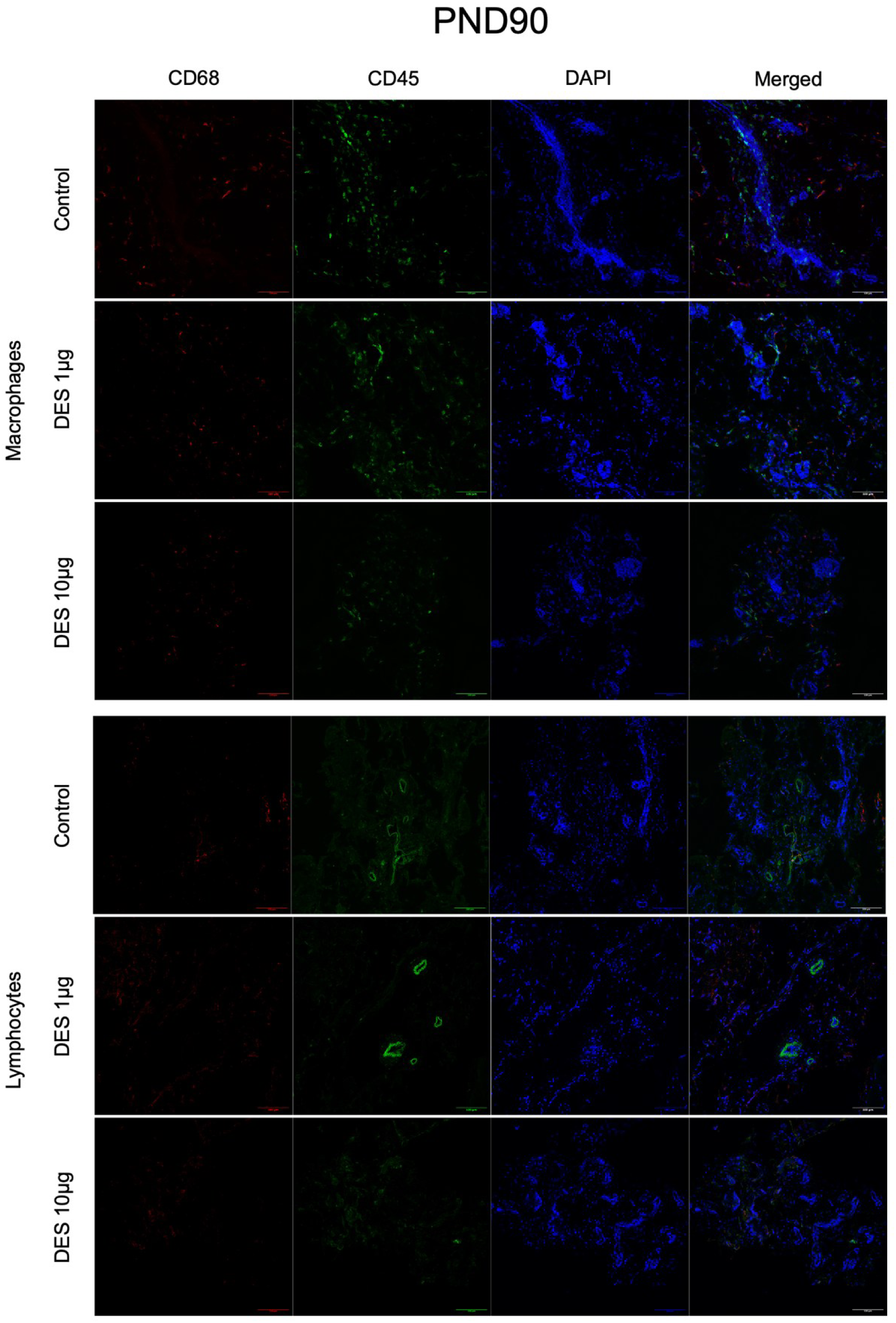
Immune cell quantification at PND90 after DES exposure in the mammary gland. Immunofluorescence staining using markers CD45 and CD68 for macrophages (**A**) and CD45 and CD3 for lymphocytes (**B**). Magnification 10x, scale bar 300 μm.

